# Impaired migration and metastatic spread of human melanoma by a novel small molecule targeting the transmembrane domain of death receptor p75^NTR^

**DOI:** 10.1101/2023.11.13.566904

**Authors:** Vanessa Lopes-Rodrigues, Samuel A. Nyantakyi, Xueqing Lun, Jianbo Zhang, Ajeena Ramanujuan, Shuhailah Salim, Michael Saleeb, Donna L. Senger, Carlos F. Ibáñez

## Abstract

Receptor transmembrane domains (TMDs) are crucially involved in relaying ligand information from extracellular to intracellular spaces and represent attractive targets for small molecule manipulation of receptor function. Screening a library of over 8,000 drug-like compounds with an assay based on the TMD of death receptor p75^NTR^, we identified a novel small molecule capable of inhibiting p75^NTR^-mediated migration of human melanoma cells. Employing medicinal chemistry, a more potent derivative termed Np75-4A22 was identified that blocks nerve growth factor (NGF)-mediated melanoma invasion at submicromolar concentrations. Mechanistically, Np75-4A22 was found, at least in part, to function by antagonizing NGF-mediated recruitment of the actin-bundling protein fascin to p75^NTR^ and its association with the actin cytoskeleton. Importantly, preclinical assessment of Np75-4A22 showed high oral bioavailability, low toxicity, and significant inhibition of melanoma lung metastases in a highly metastatic mouse model. These results support further development of this approach as an alternative or complementary strategy for patients that do not respond to conventional chemotherapy or immune checkpoint inhibitors.

## Introduction

Melanoma is the most serious form of skin cancer, and its worldwide incidence has been rising rapidly ^1^. In 2020, there were an estimated 325,000 new cases of melanoma worldwide, accounting for 1.7% of all global cancer diagnoses and 57,000 deaths ^2^. This high mortality rate has been associated with the highly invasive and metastatic nature of melanoma ^2^. In patients with metastatic melanoma carrying mutated forms of BRAF, which comprise about 40 to 50% of all melanoma patients ^3^, tumor regression and prolonged overall survival can be achieved with vemurafenib and dabrafenib, two selective BRAF inhibitors ^4^. However, resistance and disease progression has been observed in virtually every patient treated with these inhibitors ^4^. Strategies that target melanoma with mutant KIT have been effective however activating mutations in KIT are only present in a small percentage of melanoma patients ^5^. Thus, over 50% of melanoma lack a clear target for therapeutic intervention, and in those that do, acquired resistance is an ensuing problem. Currently, immune checkpoint inhibitors (ICI) are a primary treatment option available to patients with metastatic melanoma, and while these treatments provide long-term survival in some patients, recent clinical trials with ICIs indicate that over 50% of metastatic melanoma patients fail to respond to mono-therapy with either Nivolumab or Ipilimumab (targeting PD-1 and CTAL-4, respectively) ^6^ and objective or complete response rates are in the order of 60% and 20% respectively ^6,7^ when used in combination. Why almost 40% of these patients do not respond to the very best combination of available immune therapy is not understood.

The p75 neurotrophin receptor (p75^NTR^, also known as NGFR, TNFRSF16 and CD271) is a member of the death receptor superfamily, characterized by the presence of a 6-helix globular domain in the intracellular region known as the “death domain” ^8–10^. Prototypical death receptors include the Tumor Necrosis Factor Receptor 1 (TNFR1) and the Fas receptor (CD95). In addition to inducing apoptotic cell death in subpopulations of neurons, glia, and cancer cells ^11–13^, p75^NTR^ has been shown to regulate cell motility and migration in response to neurotrophins, such as nerve growth factor (NGF) ^14–16^ and is expressed at high levels in the neural crest and its cellular derivatives, including melanocytes ^17,18^. p75^NTR^ is present in deeply invasive melanoma lesions ^15^ and has been associated with tumor cell plasticity ^19–21^, progression ^22^, and metastases ^22,23^. Numerous studies have highlighted the ability of p75^NTR^ to regulate the survival and migration of melanoma cells in response to neurotrophins^15,21,24–28^ and have implicated p75^NTR^ in resistance to targeted BRAF and MEK inhibition ^29,30^ and immune based therapies ^31–33^. Together, these studies highlight the functional role of p75^NTR^ and its potential as a therapeutic target for interventions aimed at mitigating metastatic melanoma.

The mechanism by which p75^NTR^ becomes activated by endogenous neurotrophin ligands involves a conformational rearrangement of pre-formed receptor dimers that is transferred from extracellular to intracellular domains through a highly conserved transmembrane Cys residue (Cys^256^ in human p75^NTR^) ^34^. This Cys residue forms an intramembrane disulphide bridge that stabilizes constitutive dimers at the plasma membrane ^34^. Neurotrophin binding to p75^NTR^ induces a conformational change measurable by fluorescence resonance energy transfer (FRET) anisotropy that results in rapid but transient separation of receptor intracellular domains in an oscillatory fashion. Replacement of the transmembrane Cys to Alanine abolishes this conformational change and blunts downstream signaling in response to neurotrophins without affecting receptor expression at the cell surface. Knock-in mice carrying this replacement show significant protection from epileptic-induced neuronal damage ^35^ as well as neuropathology and memory impairment caused by over-expression of the 5xFAD Alzheimer’s disease transgene ^36^. The crucial role played by TMD-TMD interactions in the mechanism of p75^NTR^ activation suggests that targeting the receptor TMDs may represent a novel approach to develop small molecule modulators of p75^NTR^. Interestingly, the p75^NTR^ TMD is fairly unique: the highest scoring protein in the mammalian genome, olfactory receptor 52E4, has less than 30% sequence identity in this region, supporting this TMD as a suitable drug target. In a proof-of-principle study using a library of small molecules, our laboratory adapted a screening assay based on the AraTM transcription factor system in bacteria ^37,38^ to identify a chalcone flavonoid, NSC49652, that interacted directly with the p75^NTR^ TMD and induced profound conformational changes and activity in the full-length receptor in mammalian cells ^39^. The compound triggered apoptotic cell death that was dependent on p75^NTR^ and JNK activity in neurons and melanoma cells, and inhibited tumor growth in a melanoma mouse model ^39^.

In the present study, we used the AraTM assay to screen a much larger and chemically diverse library of over 8,000 compounds with drug-like properties for small molecules targeting the p75^NTR^ TMD and capable of affecting receptor function. We report the discovery of a novel molecule that was improved by medicinal chemistry to generate a derivative with good oral bioavailability, low toxicity, and the ability to block melanoma invasion *in vitro* and metastases *in vivo*.

## Results

### A novel pyrazine interacts with the p75^NTR^ TMD and induces dynamic changes in the full-length receptor in mammalian cells

The AraTM assay ^37,38^ was used to screen a library of approximately 8,000 compounds from the Chemical Biology Consortium Sweden (CBCS) for molecules capable of interacting with the p75^NTR^ TMD as previously described ^39^. Unlike our previous pilot study, which used a small collection (≈1,500 compounds) of known drugs from the National Cancer Institute (USA), the CBCS library was curated to represent a much larger chemical diversity, and to contain novel compounds that have drug-like properties with respect to parameters such as molecular weight, hydrogen bond accepting and donating groups, lipophilicity, and polar surface area. Briefly, a fusion protein consisting of the maltose binding protein (MBP) at the N-terminus, followed by the p75^NTR^ TMD and the AraC bacterial transcription factor at the C-terminus was expressed in the AS19 strain of *E. coli* which lacks lipopolysaccharide (LPS) in the cell wall, thereby increasing the access of different types of drugs and macromolecules to targets in the plasma membrane and cytoplasm (Good 2000). TMD-mediated dimerization of AraC leads to its activation and induction of a GFP reporter expressed from a second plasmid carrying AraC binding sites in its promoter region. Small molecules may reduce or enhance TMD-TMD interactions, leading to decreased or increased GFP readouts, respectively. Compounds that affect bacterial growth (i.e. reduce OD630 by more than 30%) are considered toxic and removed from the screen. Compounds showing a statistically significant (*p*<0.05) change in the normalized GFP readout greater than 50% were retained in the screen and subjected to secondary and tertiary screens for reduced toxicity and increased potency and specificity (**Figure 1A**). This led to the identification of a previously undescribed pyrazine that we termed Div17E5. The molecule consists of a pyrazin-2-yl phenolic moiety joined at position 6 by a 3,4-dihalogenated anilino substituent (**Figure 1B**). In dose-response studies, Div17E5 displayed an IC_50_ of 12μM in the AraTM assay and showed good selectivity for the p75^NTR^ TMD as compared to TMDs of two unrelated proteins (**Figure 1C**). Mutation of TMD residues Ile^252^ or Val^254^ abolished the ability of Div17E5 to affect the p75^NTR^ TMD (**Figures 1D, E**), suggesting they are key residues for drug binding. On the other hand, mutation of Pro^253^ had no effect (**Figure 1F**). None of the mutations diminished baseline interaction between TMDs in the absence of drug (**Figures 1D-F**), indicating that Div17E5 interacted with p75^NTR^ TMD residues that do not normally contribute to TMD dimerization. The fact that point mutations cancelled the effects of Div17E5 in the AraTM assay suggests that the compound targeted the TMD and not the technology of the assay.

**Figure 1.**
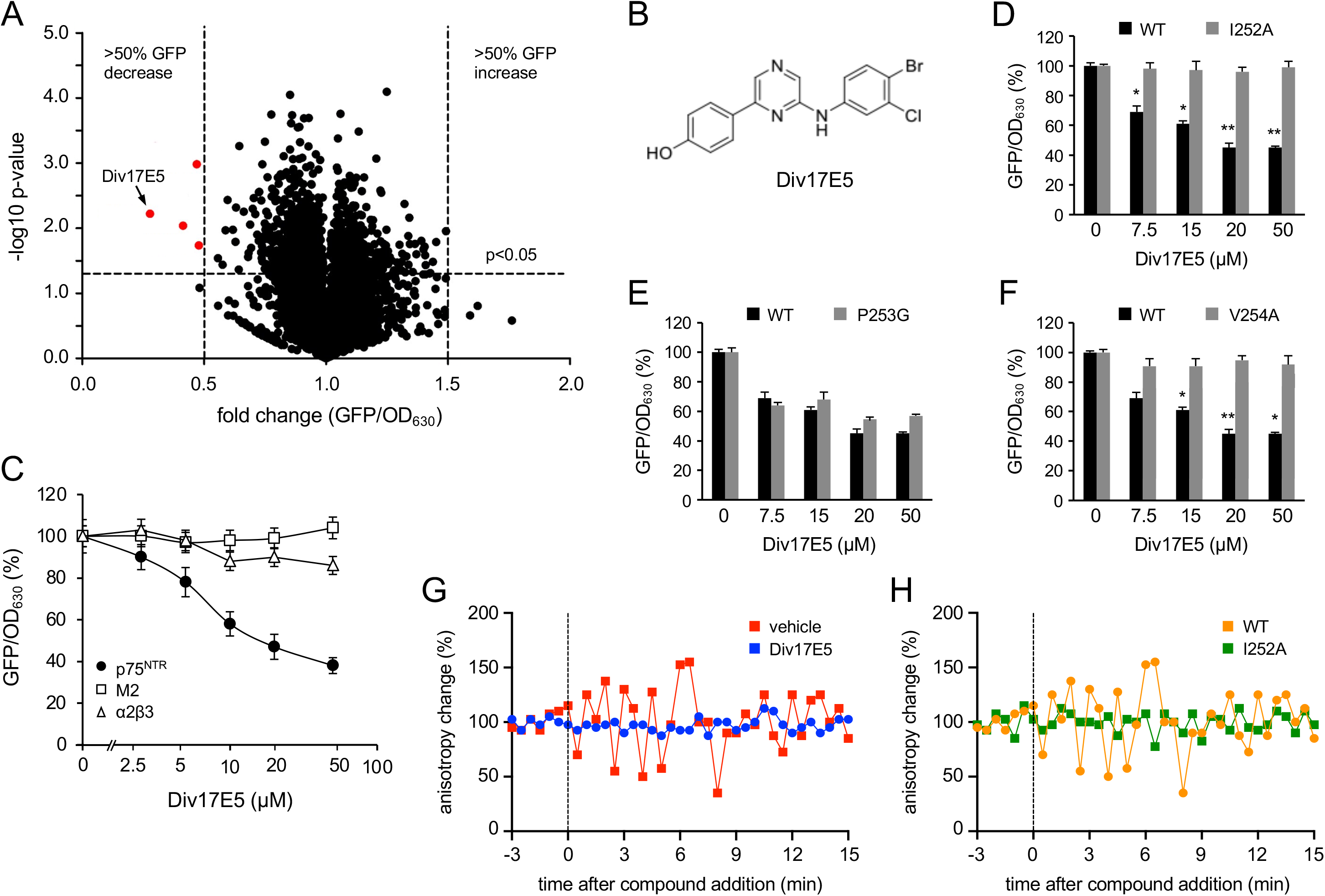
Novel pyrazine Div17E5 interacts with the p75^NTR^ TMD and induces dynamic changes in the full length receptor in mammalian cells. (A) Volcano plot of AraTM screening assay of Screening Set v2010 (8,482 compounds) from the Chemical Biology Consortium Sweden (www.cbcs.se). Fold change was calculated against vehicle (DMSO) and hits were defined as compounds resulting in greater than ±0.5 fold change in GFP/OD630 signal without affecting OD630 by greater than 0.3 fold across 3 independent runs. Compound Div17E5 is indicated. (B) Chemical structure of Div17E5. (C) Dose response of Div17E5 in the AraTM assay of p75^NTR^ TMD in comparison with unrelated TMDs from α2β3 integrin and Matrix-2 protein (M2) from the viral envelope of influenza A virus. Results are plotted as mean ± SD (N=3). (D-F) Comparison of wild type (WT) p75^NTR^ TMD and I252A, P253G and V254A mutants in the AraTM assay in response to increasing doses of Div17E5. GFP/OD_630_ signal for WT TMD without any drug was set at 100% and all other measurements are relative to that. Results are plotted as mean ± SD (N=3). *, p<0.05: **, p<0.01. (G) Live cell homo-FRET anisotropy in response to Div17E5 of full-length, wild-type human p75^NTR^ expressed in COS cells. Shown are representative traces of average anisotropy change after addition of Div17E5 (10μM) or vehicle. (H) Live cell homo-FRET anisotropy in response to Div17E5 of full-length, wild-type human p75^NTR^ in comparison to I252A p75^NTR^ TMD mutant. Shown are representative traces of average anisotropy change after addition of Div17E5 (10μM).

Previous real-time homo-FRET anisotropy studies by ours as well as other laboratories have shown that the intracellular death domains of p75^NTR^ are associated with each other (high FRET, low anisotropy state) under basal conditions ^34,40^, but neurotrophin binding induces transient separation of p75^NTR^ death domains (low FRET, high anisotropy state) manifested as oscillations in real-time anisotropy measurements ^35,39,41^. COS cells expressing full length, GFP-tagged rat p75^NTR^ were treated with vehicle or Div17E5 and changes in anisotropy levels were recorded over time. The compound induced oscillations in real-time anisotropy (**Figure 1G**), similar to those induced by endogenous p75^NTR^ ligands, such as NGF ^35,39,41^. In agreement with the AraTM data, mutation of transmembrane residue Ile^253^ (equivalent to human Ile^252^) to Ala blunted homo-FRET changes induced by Div17E5 (**Figure 1H**). Together, these findings suggest that Div17E5 binding to the TMD of p75^NTR^ induces conformational changes in the full-length receptor that can be propagated to intracellular death domains.

### Div17E5 induces apoptosis of human melanoma cells in a p75^NTR^-dependent manner

One of the main biological readouts of p75^NTR^ activity in response to neurotrophins is the induction of caspase-mediated apoptotic cell death ^11–13^. We investigated whether Div17E5 displayed p75^NTR^-mediated pro-apoptotic activity in the human melanoma cell line A875, which expresses high levels of this receptor ^42^, and a derived line in which p75^NTR^ expression had been abolished by shRNA knock-down (A875-NT and A875-shp75, respectively, **Figure 2A**). Div17E5 treatment induced apoptosis, as indicated by cleavage of Poly (ADP-ribose) Polymerase (PARP), in control A875-NT cells, while p75^NTR^ knock-down significantly reduced the pro-apoptotic effects of the compound (**Figure 2B**). Apoptosis induced by Div17E5 could be blocked by the pan-caspase inhibitor Q-VD-pOH (**Figure 2C**). In line with these findings, Div17E5 decreased viability of A875 melanoma cells in a dose-dependent manner, while A875 cells lacking p75^NTR^ were largely refractory to the compound at doses up to 10μM (**Figure 2D**). Above this concentration, both cell lines were sensitive to the compound, indicating *in vitro* off-target effects at high concentrations, which is not unusual for target specific drugs.

**Figure 2.**
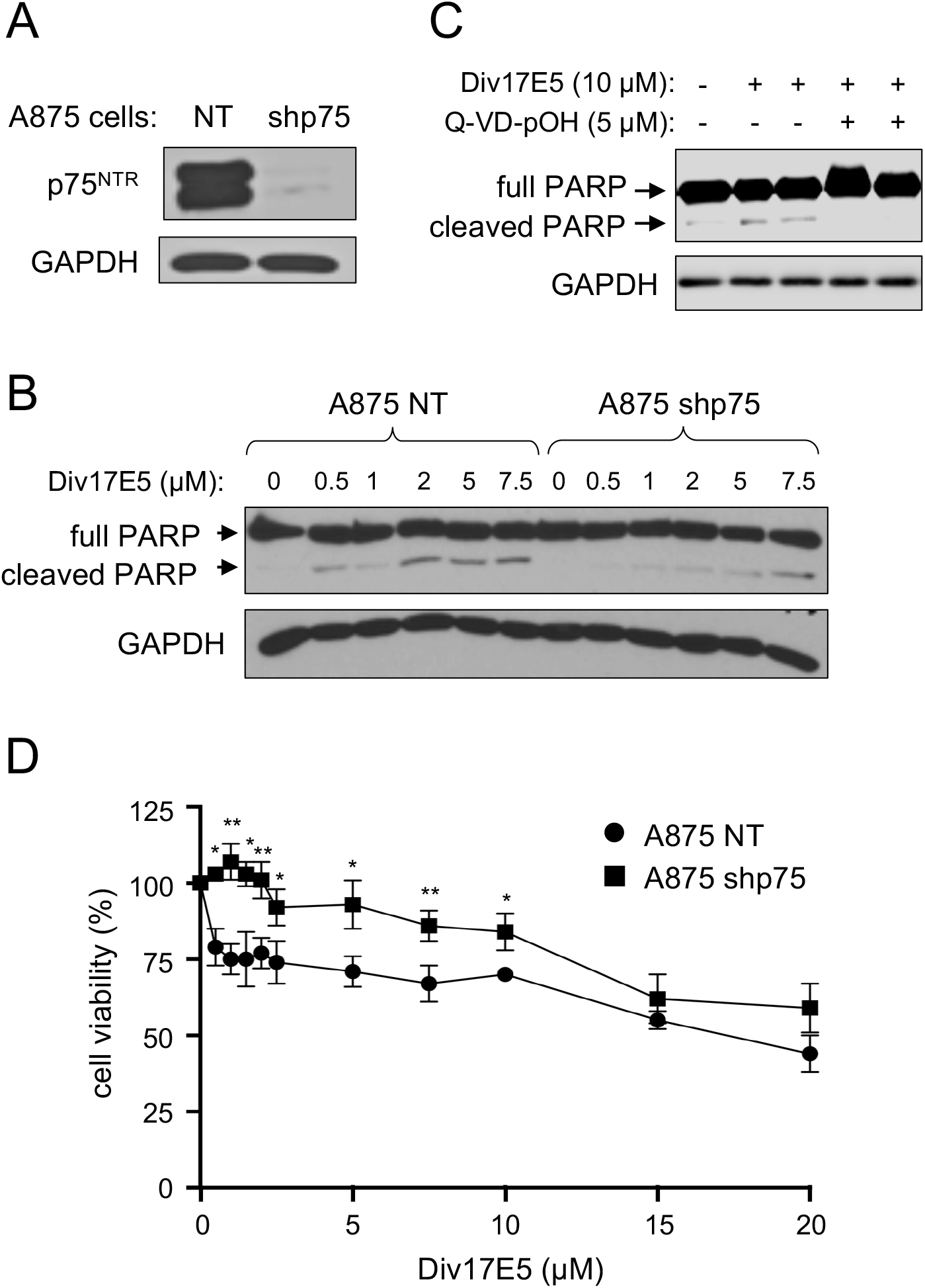
Div17E5 induces apoptosis of human melanoma cells in a p75^NTR^-dependent manner. (A) Western blot analysis of cleaved PARP in A875 melanoma cells in response to Div17E5 (10µM) in the presence or absence of pan-caspase inhibitor Q-VD-pOH (5µM). Reprobing for GAPDH was used as loading control. The experiment was repeated three times with comparable results. (B) Expression of p75^NTR^ in A875 control (NT, non-targeting) and knock-down (shp75) cells. (C) Dose response analysis of cleaved PARP induction by Div17E5 in A875 control (NT) and knock-down (shp75) cells. (D) Dose-dependent cell viability of A875 control (NT) and knock-down (shp75) cells in response to Div17E5. Results are plotted as mean ± SD (N=3). *, p<0.05; **, p<0.01.

### Div17E5 impairs cell motility and chemotaxis of human melanoma cells through targeting p75^NTR^ but independent of its pro-apoptotic effects

As p75^NTR^ is known to regulate motility and migration in melanoma cells, we tested whether Div17E5 affected these activities. Using the wound-healing assay in cell monolayers, we found that Div17E5 significantly impaired motility of A875-NT cells (**Figures 3A, B**). Importantly, the effect of Div17E5 on cell motility was not due to apoptosis, as it could not be rescued by treatment with the caspase inhibitor Q-VD-pOH (**Figures 3A, B**). Div17E5 did not significantly affect wound closure in A875-shp75 cells (**Figure 3C**), suggesting that its effects on cell motility are mediated through p75^NTR^.

**Figure 3.**
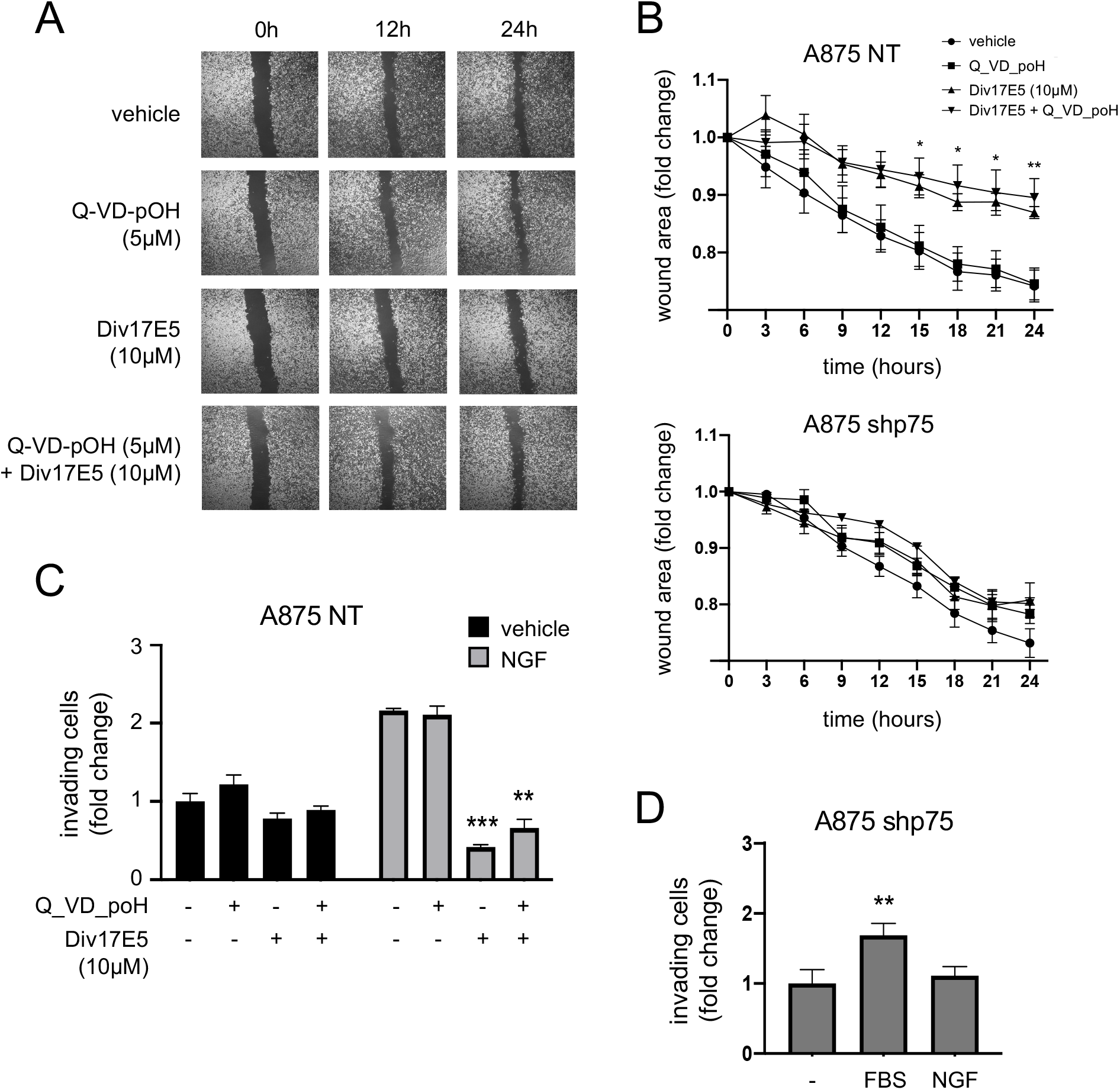
Div17E5 impairs cell motility and chemotaxis of A875 melanoma cells through p75^NTR^. (A) Phase contrast photomicrographs of wound healing cell motility assay in A875 cells treated with vehicle or Div17E5 in the presence or absence of pan-caspase inhibitor Q-VD-pOH. Size bar, Xmm. (B, C) Time course of wound healing in A875-NTcontrol (B) and knock-down shp75 (C) cells in response to Div17E5 in the presence or absence of Q-VD-pOH. Results are plotted as mean ± SD (N=3). *, p<0.05; **, p<0.01. (D) Boyden chamber analysis of A875 cell chemotaxis in response to NGF in the presence or absence or Div17E5 or pan-caspase inhibitor Q-VD-pOH. Results are plotted as mean ± SD (N=3). **, p<0.01; ***, p<0.001. (E) Boyden chamber analysis of cell chemotaxis of knock-down (shp75) A875 cells in response to NGF or serum (FBS). Results are plotted as mean ± SD (N=3). **, p<0.01.

These results led us to hypothesize that Div17E5 may antagonize NGF-induced chemotaxis of A875 melanoma cells, which has previously been shown to be mediated by p75^NTR^ ^25,43^. We tested this in trans-well Boyden chambers with NGF in the lower compartment by assessing the extent of cell migration across the membrane in the presence and absence of DivE175 and caspase inhibitor. A875-NT cells responded robustly to NGF, while the knock-down cells did not, although they still responded to serum (**Figure 3D, E**). Div17E5 did not significantly affect background cell migration in the absence of NGF but strongly inhibited NGF-induced chemotaxis of A875-NT cells (**Figure 3D**). This effect could not be rescued by the caspase inhibitor, indicating that it was not due to apoptosis (**Figure 3D**). Together these results demonstrate that Div17E5 can interfere with cell motility and chemotaxis of human A875 melanoma cells through p75^NTR^.

### Medicinal chemistry defines structural requirements of Div17E5 activity and identifies a more potent analog

We applied medicinal chemistry to investigate the molecular determinants underlying Div17E5 activity and to identify more potent and better tolerated compounds. We synthesized 31 chemically diverse analogs (Series 1) carrying different groups linked to either of the two benzene rings (**Figure 4A**). R1 and R2 groups ranged from electron-donating groups, electron-withdrawing groups, and bulky hydrophobic groups in different combinations, all the while maintaining the -NH-linker at position 6 (Series 1, compounds MS_11 to MS_71, MS_100 and MS_114 in **Supplementary Figure 1A**). To understand the essence of the amino (-NH-) linker (**Figure 4B**), 6 additional analogs lacking it were also synthesized (Series 2A, MS_72 to MS_78 in **Supplementary Figure 1A**). Finally, a carbon linker was introduced at position 2 of the pyrazine in MS_63 (**Figure 4C**) to generate an additional 4 compounds (Series 2B, MS_109 to MS_112 in **Supplementary Figure 1A**). All 6 compounds from Series 2A were inactive (not shown), stressing the importance of the amino group for the activity of Div17E5. Neither did the 4 compounds in Series 2B result in improved activity (not shown). From these studies, only two analogs from Series 1 were retained, namely MS_18 and MS_63, which displayed similar IC_50_s as Div17E5 in the AraTM assay but with somewhat lower bacterial toxicity, as assessed by OD_630_ measurements (not shown). MS_18 and MS_63 featured electron-donating 4-methoxy and 3-dimethylamino, respectively, as the R1 group, and electron-withdrawing 4-nitro and 4-bromo-2,3-dichloro motifs, respectively, as the R2 group. However, when tested in the wound healing assay in A875-NT cells, both compounds were less potent than Div17E5 (**Figure 4D**).

**Figure 4.**
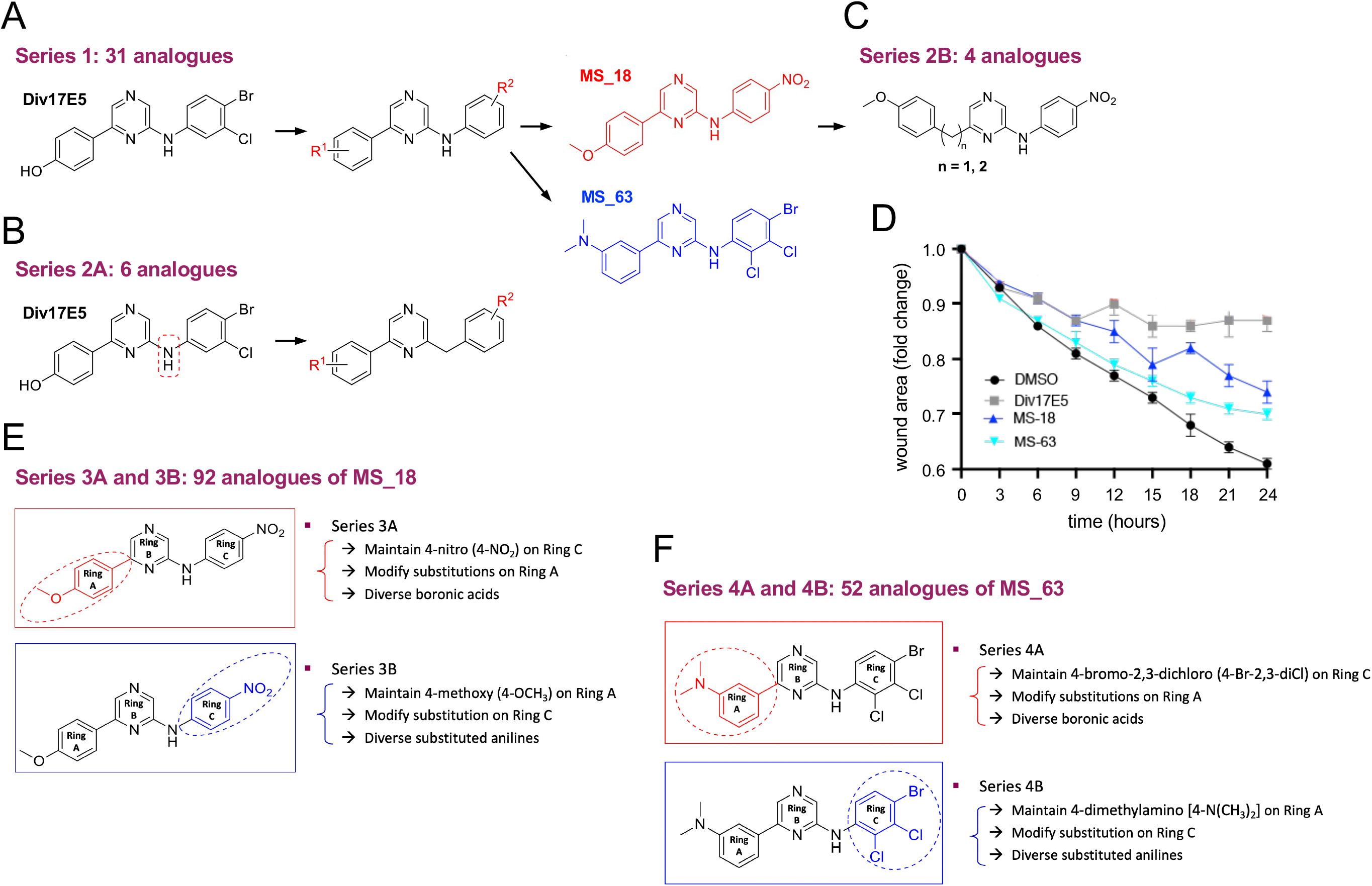
Medicinal chemistry strategy for derivation of Div17E5 analogs. (A) Series 1 included 31 chemically diverse analogs carrying different groups linked to either of the two benzene rings (R1 and R2). Chemical structures of MS_18 and MS_63, the two compounds retained from this series, are shown. (B) Series 2B included 6 analogs lacking the amino (-NH-) linker with. R1 and R2 groups. (C) Series 2C included 4 analogs carrying an extra carbon linker at position 2 of the pyrazine in MS_63. (D) Time course of wound healing in A875 NT cells in response to 10µM Div17E5, MS_18 or MS_63. Results are plotted as mean ± SD (N=3). (E) Series 3A and 3B included 92 analogs of MS_18 that maintained the 4-nitro (4-NO2) substituent on ring C while modifying substitutions on ring A (3A), or the 4-methoxy (4-OCH3) substituent on ring A while modifying substitutions on ring C (3B). (F) Series 4A and 4B included 52 analogs of MS_63 using the same principles as Series 3A and 3B, respectively.

MS_18 and MS_63 were then further modified to yield Series 3A/B and 4A/B, respectively. The design strategy for analogs in Series 3A was to maintain the 4-nitro (4-NO2) substituent on ring C found in MS_18, while modifying substitutions on ring A (**Figure 4E**). Conversely, for 3B, we maintained the 4-methoxy (4-OCH3) substituent on ring A found in MS_18, while modifying substitutions on ring C. The same design principles were employed for Series 4A and 4B on MS_63 (**Figure 4F**). In total, 144 new compounds were synthesized and tested in the AraTM assay. A detailed analysis of structure-activity relationships for compounds in Series 3A/B and 4A/B is presented separately (**Supplementary Text**). From these efforts, Np75-4A22 was retained (**Figure 5A**), as it showed better performance at low micromolar concentrations and improved IC_50_ compared to Div17E5 (**Figure 5B**). The activity of this compound in the AraTM assay was further characterized using mutant p75^NTR^ TMDs. Similar to Div17E5, Np75-4A22 was affected by replacement of Ile^252^ and unaffected by mutation of Pro^2^^53^ (**Figures 5C, D**). However, unlike Div17E5, Np75-4A22 was unaffected by mutation of Val^2^^54^ (**Figure 5E**), indicating that the two compounds interact with a partially overlapping but distinct set of residues in the p75^NTR^ TMD.

**Figure 5.**
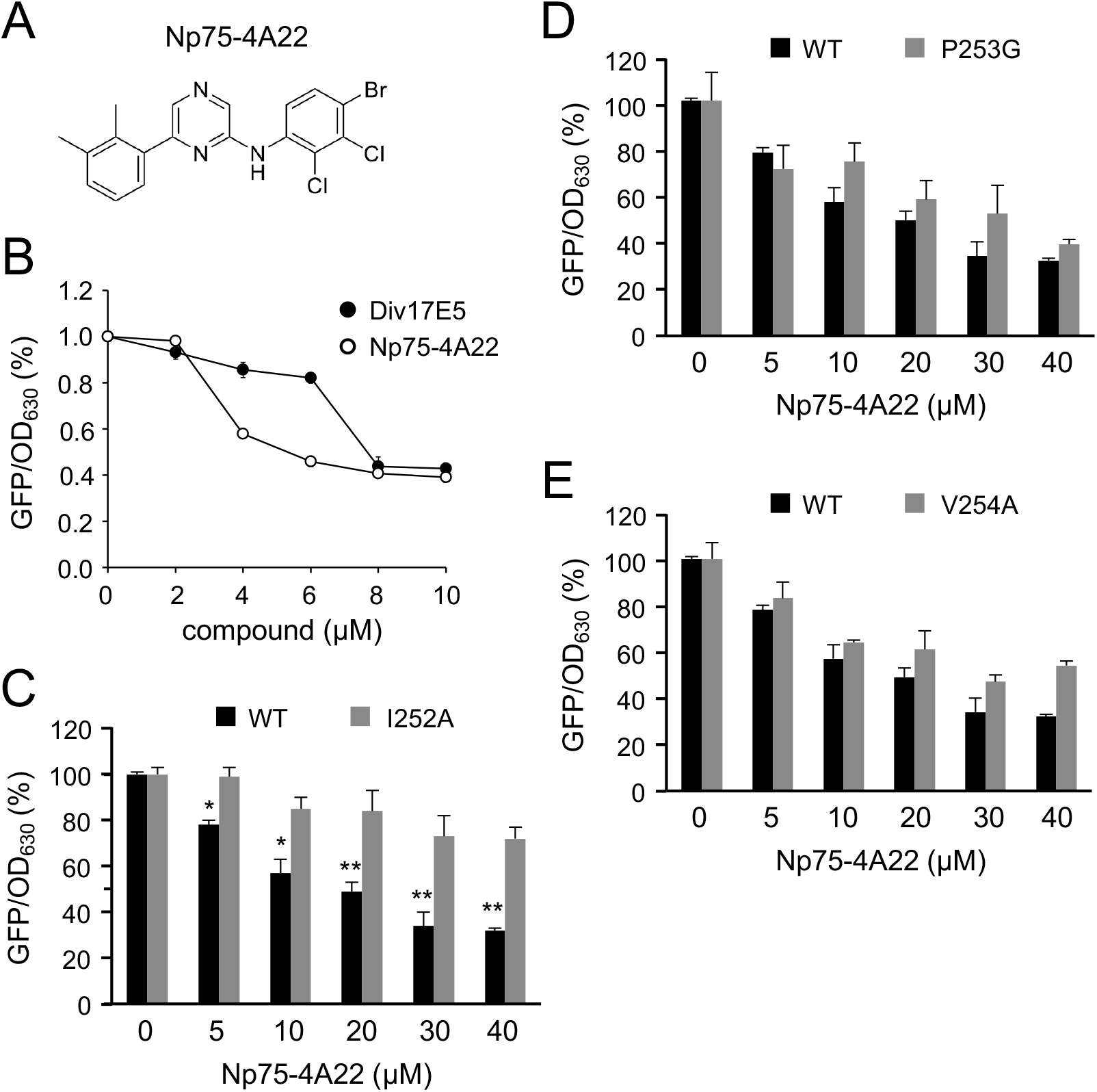
Np75-4A22, a more potent analog of Div17E5 identified by medicinal chemistry. (A) Chemical structure of Np75-4A22. (B) Dose response of Div17E5 and Np75-4A22 in the AraTM assay of p75^NTR^ TMD. Results are plotted as mean ± SD (N=3). **, p<0.01. (C-E) Comparison of wild type p75^NTR^ TMD and I252A, P253G and V254A mutants in the AraTM assay in response to increasing doses of Np75-4A22. GFP/OD_630_ signal without any drug added was set at 100%. Results are plotted as mean ± SD (N=3). *, p<0.05: **, p<0.01.

### Np75-4A22 inhibits melanoma cell invasion at submicromolar concentrations without inducing cell death

In contrast to Div17E5, Np75-4A22 did not induce apoptosis of A875 melanoma cells in the submicromolar and low micromolar range, as assessed by PARP cleavage (**Figure 6A**). However, Np75-4A22 was significantly more potent than Div17E5 in the wound healing assay, displaying considerable inhibitory activity at low micromolar and submicromolar concentrations, at which Div17E5 was totally inactive (**Figure 6B**). In the Boyden chamber assay of cell chemotaxis, 1μM Np75-4A22 blocked all NGF activity, while Div17E5 was inactive at that concentration (**Figure 6C**). Interestingly, Np75-4A22 was also capable of significantly reducing NGF-induced A875 chemotaxis at submicromolar concentrations (**Figure 6D**). Thus, the Np75-4A22 analog was less toxic to melanoma cells but considerably more potent at blocking melanoma cell invasion than Div17E5.

**Figure 6.**
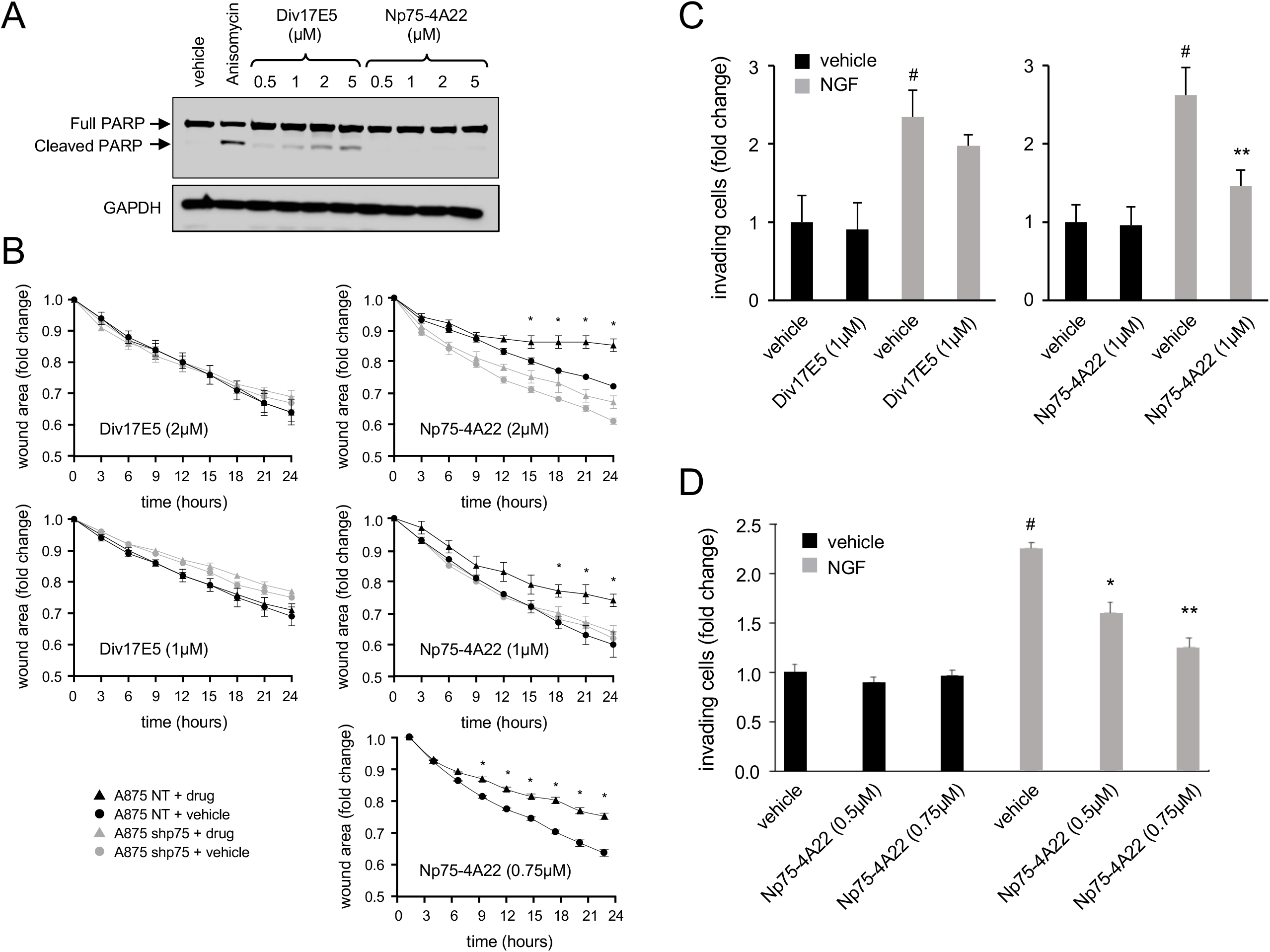
Np75-4A22 inhibits melanoma cell invasion at submicromolar concentrations without inducing cell death. (A) Western blot analysis of cleaved PARP in A875 melanoma cells in response to increasing concentrations (0.5 to 5µM) of Div17E5 and Np75-4A22. A potent inducer of apoptosis, anisomycin (1uM), was used as positive control. Reprobing for GAPDH was used as loading control. The experiment was repeated three times with comparable results. (B) Time course of wound healing in A875 control (NT) and knock-down (shp75) cells in response to low micromolar and submicromolar concentrations of Div17E5 (left side) or Np75-4A22 (right side). Results are plotted as mean ± SD (N=3). *, p<0.05. (C) Boyden chamber analysis of A875 cell chemotaxis in response to NGF in the presence or absence or Div17E5 (black bars) or Np75-4A22 (gray bars). Results are plotted as mean ± SD (N=3). #, p<0.05 vs. no NGF; *, p<0.05 vs. no drug. (D) Boyden chamber analysis of A875 cell chemotaxis in response to NGF in the presence of submicromolar concentrations of Np75-4A22. Results are plotted as mean ± SD (N=3). #, p<0.05 vs. no NGF; *, p<0.05 and **, p<0.01 vs. no drug.

### Np75-4A22 blocks NGF-mediated recruitment of fascin to p75^NTR^ and fascin association with the cytoskeleton

It has been shown that the actin-bundling protein fascin interacts with p75^NTR^ in an NGF-dependent manner ^15^. NGF also enhanced the association of fascin with the actin cytoskeleton in melanoma cells, as shown by increased fascin levels in detergent-insoluble fractions upon NGF treatment ^15^. Importantly, the same study also showed that NGF-induced migration of A875 melanoma cells was inhibited by a mutant of fascin that precludes its binding to actin. Together, these findings indicated that one mechanism by which NGF enhances migration of melanoma cells is through fascin recruitment to p75^NTR^ as well as increased fascin association with the actin cytoskeleton. We investigated whether Np75-4A22 has any effects on the ability of NGF to regulate fascin in A875 melanoma cells. Fascin co-immunoprecipitated with p75^NTR^ in control NT cells but not in shp75 knockdown cells (**Figure 7A**). In agreement with the previous study, NGF treatment strongly stimulated the association of fascin with p75^NTR^ (**Figure 7A**). Interestingly, treatment with Np75-4A22 at 1μM completely suppressed the effect of NGF reducing fascin/p75^NTR^ interaction to baseline levels (**Figure 7A**). Moreover, while NGF increased the levels of fascin in the detergent insoluble fraction of A875-NT cells, 1μM Np75-4A22 also blocked this effect (**Figure 7B**). These results indicated that, mechanistically, Np75-4A22 may inhibit melanoma cell invasion by interfering with NGF-mediated recruitment of fascin to the receptor and fascin association with the actin cytoskeleton. Together, our *in vitro* studies prompted us to investigate whether Np75-4A22 may be able to impair melanoma metastasis *in vivo*.

**Figure 7.**
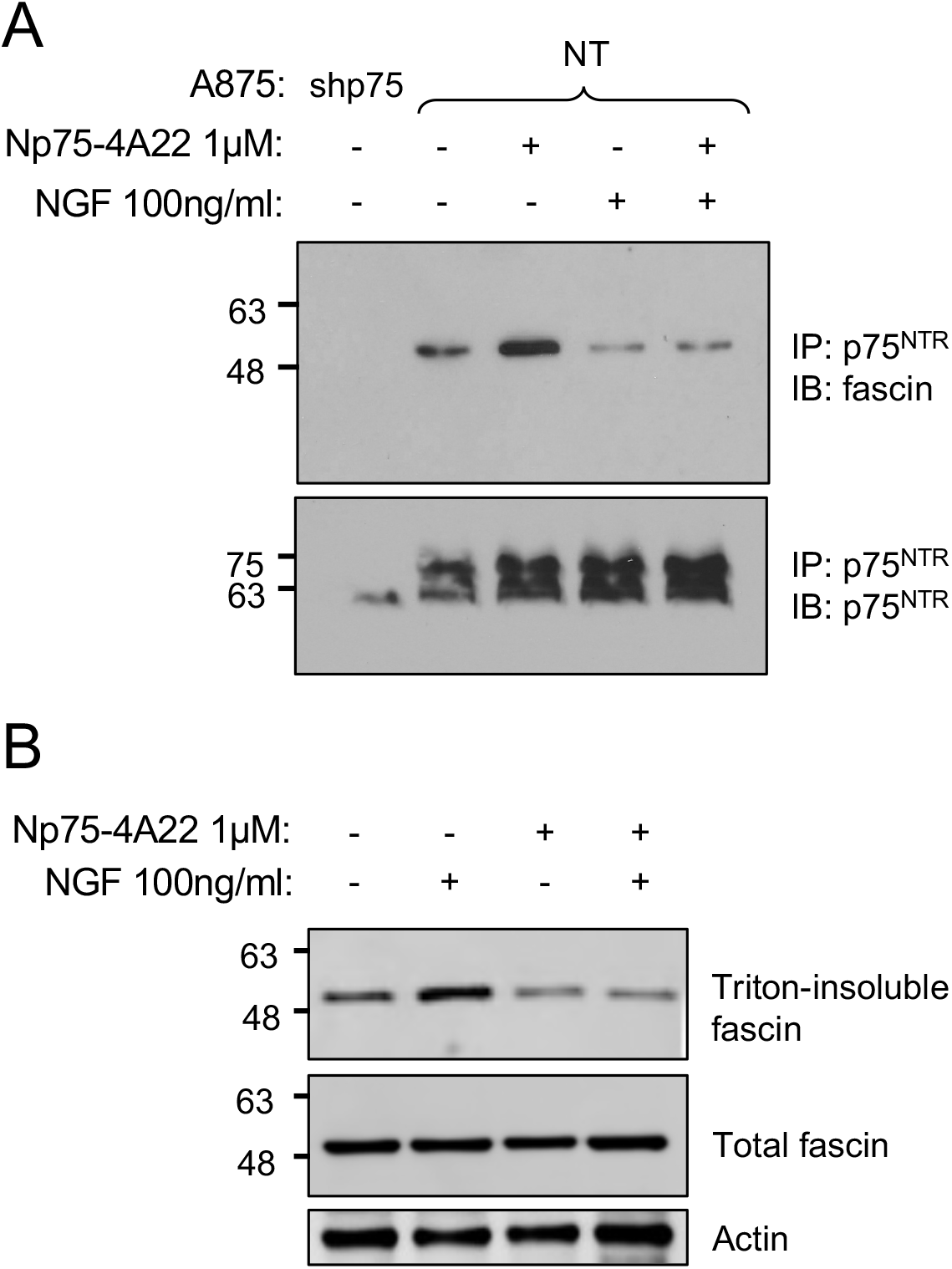
Np75-4A22 blocks NGF-mediated recruitment of fascin to p75^NTR^ and fascin association with the cytoskeleton. (A) Western blot analysis (IB) of fascin in p75^NTR^ immunoprecipitates (IP) of A875 melanoma cells after overnight treatment with NGF or Np75-4A22 as indicated. The experiment was repeated three times with comparable results. (B) Western blot analysis of fascin in Triton-insoluble extracts of A875 melanoma cells after overnight treatment with NGF or Np75-4A22 as indicated. The experiment was repeated three times with comparable results.

### Oral administration of Np75-4A22 reduces melanoma lung metastasis in a mouse xenograft model

To determine optimal dosing of Np75-4A22, a pharmacokinetic study was performed following single intravenous (3mg/kg) or oral gavage (10mg/kg) administration in female and male C57Bl/6 mice. Plasma concentration-time profiles and main pharmacokinetic parameters are summarized in Supplementary Figures S2A and B, respectively. Briefly, Np75-4A22 exhibited low clearance and long half-life following intravenous (i.v.) administration, as well as rapid absorption and good oral bioavailability following oral administration in both male and female mice. To establish maximal dosing for *in vivo* studies, toxicology analyses were performed on female and male C57Bl/6 mice 7 days after a single dose oral administration of Np75-4A22 at 50, 100 or 200mg/kg. No mortality or abnormal clinical signs were observed or treatment-related effects on body weight, food consumption or gross pathology in either male or female C57BL/6 mice treated with up to 200 mg/kg Np75-4A22. Subsequently, a repeated dose tolerability study was performed to determine toxicity of Np75-4A22 after daily oral administration at 200 mg/kg/day to female and male C57Bl/6 mice for a period of 21 consecutive days. No mortality was observed in either females or males, and no treatment-related effects were observed in males, while a mild decrease in lymphocyte cellularity and increase granulopoiesis were observed in females. In addition, and prior to *in vivo* preclinical melanoma studies, Np75-4A22 was administered by oral gavage at 200mg/kg daily (5 days on, 2 days off) for 3 weeks in immunodeficient female SCID mice. No mortality or adverse clinical signs, including body weight, food consumption or effects on gross histological pathology assessed on formalin fixed paraffin embedded lung, liver, kidney or brain were observed in any of the Np75-4A22 treated mice.

Next, to assess the potential inhibitory effects of Np75-4A22 on the development of melanoma metastases we utilized the highly metastatic human A875 xenograft model. Briefly, A875-NT or A875-shp75 cells expressing GFP-Luc (for live-monitoring of tumor growth) were injected intravenously in female SCID mice 24 hours following a single oral administration of Np75-A422 (200mg/kg) or drug vehicle alone. Mice received 3 cycles of Np75-A422 (200 mg/kg/day) following a 5 day on and 2 days off drug schedule for a total of 3 weeks (**Figure 8A**). Metastatic spread to the lungs was monitored weekly by luciferase activity using the Xenogen IVIS-200 Optical *in vivo* luciferase imaging system and mice were sacrificed after the third week. Importantly, treatment with Np75-4A22 resulted in a dramatic reduction of lung metastatic tumors in the mice that received A875 melanoma cells expressing p75^NTR^ (A875-NT; **Figure 8B**). In addition, assessment of the metastatic spread was also quantified by histologic assessment of non-consecutive lung sections stained for human nucleolin to detect the tumor cells. As shown in **Figure 8C**, a marked reduction of p75^NTR^-positive lung metastases was observed in mice treated with Np75-4A22 compared to vehicle alone. In addition, and as expected, high expression of p75^NTR^ was detected in the lung metastases established by A875-NT cells but was negligible in tumors derived from A875-shp75 cells (**Supplementary Figure S3**). Consistent with the requirement of p75^NTR^ for melanoma migration and invasion, we found that the loss of p75^NTR^ expression in A875 melanoma cells by shRNA (A875-shp75) results in a significant decrease in melanoma metastasis when compared to p75^NTR^-positive A875 (A875-NT) (**Figures 8C**), a finding consistent with previous studies on the effects of p75^NTR^ in melanoma cell dispersion (Iwamoto 1996). Together, these results show that treatment *in vivo* with Np75-4A22 can substantially decrease the spread of melanoma lung metastasis in mice. Further, we found that p75^NTR^-negative tumors (A875-shp75), although fewer and smaller than those generated by A875-NT cells, were not significantly affected by Np75-4A22 (**Figure 8C**) thus supporting the target specificity of Np75-4A22 for p75^NTR^.

**Figure 8.**
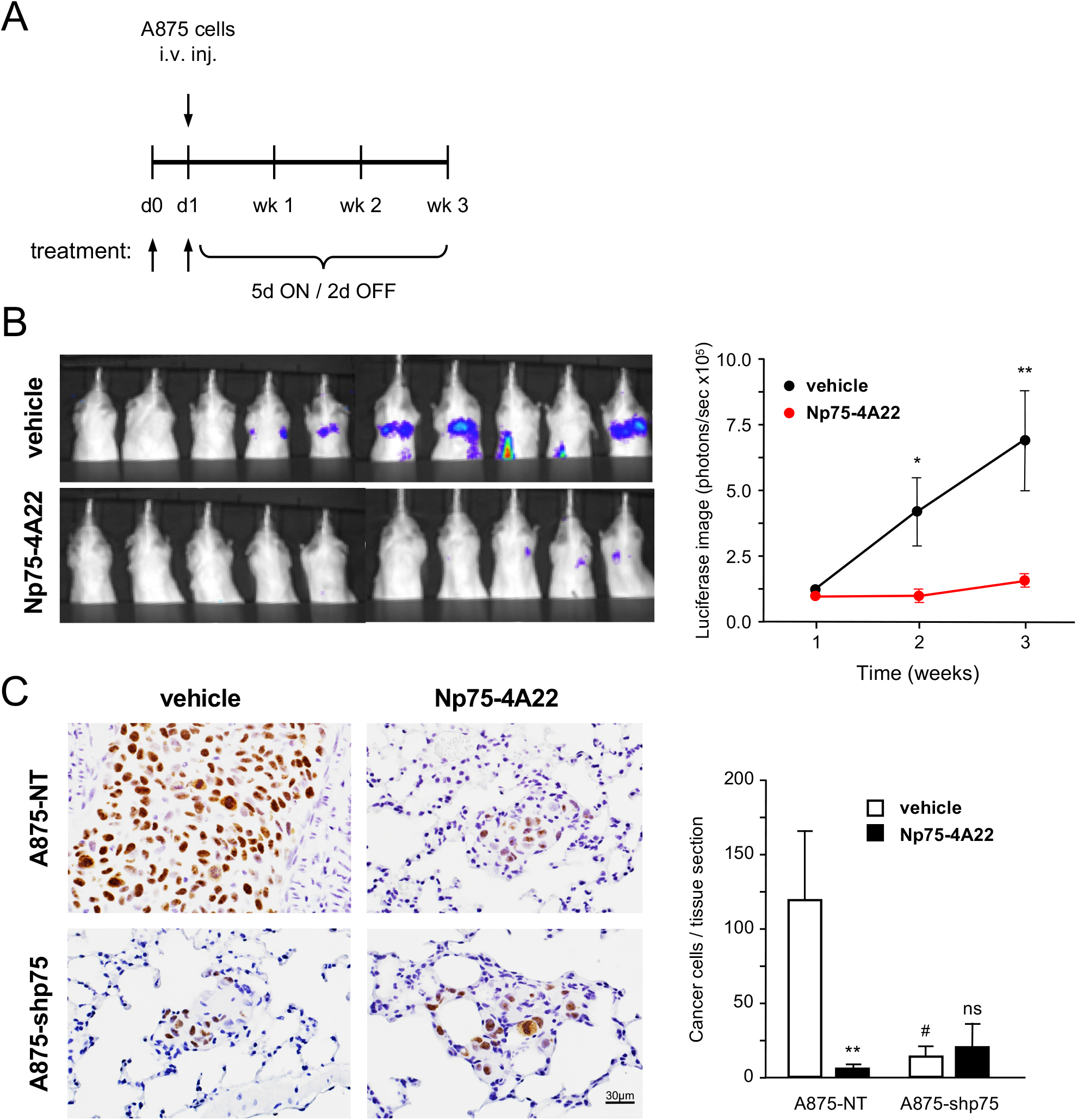
Oral administration of Np75-4A22 reduces the spread and seeding of melanoma lung metastasis in a mouse xenograft model. (A) Schematic shows the experimental design for the assessment of a human melanoma xenograft lung metastasis model following treatment with Np75-4A22. (B) Live-monitoring of lung metastatic tumor growth of A875-NT melanoma cells using the Xenogen IVIS-200 luciferase imaging system. Bioluminescent imaging of luciferase expressing A875-NT melanoma cells (control) in the lungs of SCID mice was measured 1, 2 and 3-weeks following treatment with Np75-4A22 (200 mg/kg/day) or vehicle alone. Representative images taken 3 weeks following implantation are shown (left panel). Graph shows the average quantification of luciferase signal (N=8-10 mice/group; right panel). Shown is one representative experiment from N=3 independent experiments. Data are expressed as the mean ± SEM. Means that are significantly different between treatments as analyzed by Student t-test are indicated. *, p<0.05: **, p<0.01. (C) Histological analysis of lung tumor burden in SCID mice bearing A875-NT or A875-shp75 tumors 3 weeks following of treatment with Np75-4A22 or vehicle alone. Left panel shows representative images of formalin fixed paraffin embedded lung sections stained by immunohistochemistry with a human-specific nucleolin antibody (brown) to detect the presence of tumor cells. Sections were counterstained with hematoxylin (blue). Scale bar, 60μm. Right panel shows quantification of the number of cancer cells per lung tissue section. Five non-sequential lung sections, 100 μm apart, were counted and averaged for each mouse (N=8-10 mice per group). Data is expressed as the mean ± SEM. Means that are significantly different are indicated. A875-NT vehicle vs Np75-4A22 ** p=0.0179; A875-NT vs A875-shp75 # p=0.0367; A857-shp75 vehicle vs Np75-4A22 not significant (n.s.), one-way ANOVA with Tukey’s multiple comparisons test.

## Discussion

When detected early, melanoma is highly treatable, however if left undetected it can spread and metastasize to other areas in the body with organotrophic selectivity to the lungs and brain ^44^. Herein we describe the identification of a novel small molecule targeting the TMD of p75^NTR^, a receptor implicated in the development of invasive ^15^ and metastatic ^22,23^ melanoma. This compound was found to inhibit melanoma cell migration and chemotaxis, and the generation of a more potent derivative by medicinal chemistry significantly reduced melanoma metastasis to the lungs. We predict that Div17E5 and Np75-4A22 interacts directly with the p75^NTR^ TMD as effects observed in the AraTM assay are abolished by point mutations in specific TMD residues. Interestingly, while mutation of either Ile^2^^52^ and Val^2^^54^ affected Div17E5 interaction with the p75^NTR^ TMD, only mutation in the former interfered with Np75-4A22 activity, indicating that the two compounds interact with partially overlapping but distinct TMD residues. Also notable is the observation that neither compound was affected by mutation of Pro^253^, a substitution we previously showed to interfere with the activity of NSC49652, a compound identified in a previous study ^39^, again suggesting distinct modes of action of compounds targeting the p75^NTR^ TMD.

To understand structure-function relationships of the drug/TMD interaction, and to improve the activity and safety profile of Div17E5, a series of derivatives (185 in total) were synthesized and studied. Series 1 and 2 analogs were initially made to understand the preferred pharmacophore for optimal binding. Modifications ranging from electron-donating groups, electron-withdrawing groups, and bulky hydrophobic and steric groups in different combinations, all the while maintaining the amino (-NH-) linker at position 6, resulted in 41 chemically diverse analogs with varying activity and toxicity profiles. This led to identification of MS_18 and MS_63 with similar IC50s as Div17E5 but lower potency in the wound healing assay. Consistent with Div17E5, electron-donating aromatic groups were preferred at position 2 of the pyrazine, with electron-withdrawing anilino motif favorable at position 6. Further, to understand the essence of the amino linker, 6 analogs, in which this was removed, were synthesized. All 6 compounds showed no activity, confirming the importance of the amino group for TMD binding. The NH-group, which can act both as a H-bond donor and acceptor, may participate in hydrogen bonding with key amino acids (e.g., Ile^2^^52^) in the TMD binding pocket thereby contributing essential binding activity. MS_18 and MS_63 were further modified to yield 144 additional compounds in Series 3A/B and 4A/B. The best compounds identified from this collection were Np75-3A21 and Np75-4A22. Both compounds carry electron-donating groups attached to the phenyl ring at position 2 of the pyrazine core (ring A), as well as electron-withdrawing motifs were attached to the anilino substituent at position 6 (ring C). The fact that these hits were representatives of Series 3A and 4A, while none of the modifications attempted in Series 3B and 4B yielded a better alternative, indicates that electron-donating groups on ring A are crucial when coupled with electron-withdrawing motifs on ring C.

Div17E5 acts as a p75^NTR^ agonist with respect to apoptosis induction, but as an antagonist of NGF and p75^NTR^-mediated cell motility and chemotaxis. We feel this behavior can be rationalized by the way in which p75^NTR^ engages with different downstream interactors and pathways. Recent research has revealed that downstream p75^NTR^ effectors compete for binding to the receptor intracellular domain ^41,45^. The terms of such competitions are likely to be dictated by the relative abundance of downstream effectors in different cell types, the presence or absence of p75^NTR^ co-receptors, the nature of the ligand engaging the receptor (e.g mature neurotrophins, pro-neurotrophins, myelin-derived ligands, etc.), as well as their relative concentrations, binding affinities, and relative timing of interaction. Thus, a ligand that enhances p75^NTR^ coupling to the NF-kB pathway, such as the mature form of NGF, can also reduce receptor binding to RhoGDI and coupling to the RhoA pathway ^46^. Interestingly, myelin associated glycoprotein (MAG) can tilt that balance in the opposite direction in the presence of the p75^NTR^ co-receptor NgR ^47,48^. In cerebellar neurons, the mature and unprocessed (pro) forms of NGF antagonize each other in the induction of p75^NTR^-mediated cell survival vs. cell death pathways, respectively ^45^. Against this background, it is reasonable to propose that Div17E5 binding to the p75^NTR^ TMD likely induces a particular arrangement of TMDs and overall conformation dynamics that favors one downstream pathway (i.e. apoptosis) at the expense of another (i.e. cell migration). Interestingly, Np75-4A22, an analog of Div17E5, was significantly more potent than Div17E5 at blocking melanoma cell invasion, without inducing cell death, at low micromolar and submicromolar concentrations. The two compounds are likely to interact differently with the p75^NTR^ TMD, as Np75-4A22, unlike Div17E5, was not affected by mutation of Val^2^^54^, suggesting that the p75^NTR^ TMD is exquisitely sensitive to small structural alterations that can bias different downstream pathways in one or another direction. Together, these results indicate that it is possible to identify small molecules targeting the p75^NTR^ TMD which can elicit distinct and selective functional downstream outcomes.

Mechanistically, we found that Np75-4A22 inhibited NGF-mediated recruitment of fascin to p75^NTR^ as well as the association of fascin with the cytoskeleton. Fascin, an actin-bundling protein, has been shown to play a role in cell motility through its localization in filopodia and lamellipodia forming at the leading edge of motile cells in response to adhesion to different types of extracellular matrix components ^49^. Moreover, fascin overexpression in kidney epithelial cells has been shown to promote their migratory activity in Boyden chambers ^50^. Importantly, fascin levels have been found to be upregulated in human breast cancer cell lines ^51^, and tumor progression in ovarian cancer, as well as invasion in breast cancer were found to correlate with fascin expression levels ^52^. Our results indicate that one of the mechanisms by which Np75-4A22 inhibits melanoma cell invasion involves interference with NGF-mediated recruitment of fascin to p75^NTR^ and fascin partitioning in the detergent insoluble fraction of melanoma cells, a proxy of its association with actin fibers. They further suggest that fascin may in part exert its role in cancer cell motility and invasion by linking p75^NTR^ to the actin cytoskeleton.

Pharmacokinetic and toxicology studies indicated that Np75-4A22 displays rapid absorption, high oral bioavailability and low or no toxicity. Preclinical studies investigating the therapeutic efficacy of Np75-4A22 on metastatic dissemination of melanoma cells from circulation into the lung, a primary metastatic site ^44^, were then performed. Using a highly metastatic melanoma xenograft model, daily administration of Np75-4A22 was shown to significantly inhibit the establishment of metastasis to the lungs. In addition, and consistent with this observation, knockdown of p75^NTR^ in human melanoma cells resulted in a similar decrease in lung metastases with no further effect observed in the presence of Np75-4A22, supporting the specificity of the compound for p75^NTR^. These results are consistent with the *in vitro* studies showing that Np75-4A22 effectively inhibited NGF-mediated cell motility and chemotaxis of melanoma cells in a p75^NTR^-dependent manner and at low micromolar concentrations. Together these data demonstrate a role for p75^NTR^ in the development of lung metastasis and establishes Np75-4A22, as a potential therapeutic inhibitor in the systemic dissemination of melanoma.

In summary, this study provides new evidence supporting the feasibility of targeting the TMDs of single-pass transmembrane receptors for the identification of small molecules capable of modifying receptor activity in useful ways. Np75-4A22 represents a promising lead in the development of therapies against melanoma metastasis which could prove particularly beneficial for patients showing poor responsiveness to conventional chemotherapy or immune checkpoint inhibitors.

## Methods

### Chemicals and antibodies

The library of 8,442 chemically diverse compounds was obtained from the Chemical Biology Consortium Sweden (CBCS, www.cbcs.se). Div17E5 was custom synthesised by Glixx Laboratories at a purity greater than 98%. Np75-4A22 was custom synthesised by SaiLife India at a purity greater than 98%. NGF was purchased from R&D Systems. Q-vd-Oph was purchased from Sigma Aldrich. Antibody against human p75^NTR^ extracellular domain used for immunoprecipitation was obtained from Alomone Labs (#ANT-007); the one used for immunofluorescence was from Promega (#G323A). Human-specific mouse polyclonal nucleolin antibody was from Abcam (#ab136649). GAPDH, Actin and PARP antibodies were purchased from Cell Signalling technologies. Fascin antibody (#MA5-11483) was purchased from Invitrogen. Secondary antibodies used for immunofluorescence were Alexa Fluor™ 488 Goat anti-Mouse IgG (H+L) Highly Cross-Adsorbed Secondary Antibody (ThermoFisher # A-11029) and Alexa Fluor™ 568 Goat anti-Rabbit IgG (H+L) Cross-Adsorbed Secondary Antibody (ThermoFisher # A-11011).

### AraTM screening assay

The AraTM assay was used to assess conformational changes and binding strength in a pair of interacting TMDs ^53,54^. The assay was done as previously described ^39^. Briefly, p75^NTR^ TMD cDNA (encoding NLIPVYCSILAAVVVGLVAYIAFKRW) was subcloned into AraTM chimera plasmid and transformed in AS19 LPS-negative E.Coli (which allows the uptake of a variety of molecules, peptides and nuclei acids) ^55^ along with the Ara-GFP reporter plasmid. The p75^NTR^ TMD sequence used is identical in all sequenced mammalian homologs of p75^NTR^, including human and mouse. An overnight culture was diluted 1:100 in fresh LB medium and allowed to grow till reached between 0.2 and 0.5 optical density (OD) 630, after which IPTG was added to a final concentration of 1mM to induce the expression of the p75^NTR^ TMD–AraTM chimera. The bacterial culture was added to a black-rim clear bottom 96-well plates (Corning #3631) pre-plated with the compounds (final concentration of 20μM). The plates were incubated at 38°C to allow IPTG-induced expression of the TM-AraTM chimera and after 4 hours were centrifuged to pellet the bacterial cells. LB media was aspirated and replaced with Phosphate Buffered Saline (PBS), and bacteria cells were resuspended by vigorous shaking for 10 min. GFP signal was measured in each well (excitation 475nm, emission 509 nm) and bacterial density was determined by measurement of OD_630_ in a microplate plate reader (BioTek).

### Homo-FRET anisotropy microscopy

Anisotropy imaging in COS-7 cells that were transfected with a rat p75^NTR^-EGFP* fusion construct was done as previously described ^39^. EGFP* carries a A207K mutation rendering monomeric EGFP. Changes in anisotropy were expressed as fold change at each time point (every 30 seconds) in comparison to the mean of 6 time points obtained prior to addition of treatment. Images were acquired using Nikon Ti-E based live cell epi-fluorescence microscope and MetaMorph software and analyzed using MatLab from Mathworks.

### Cell culture and immunoblotting

COS7 and A875 cell lines were cultured under standard conditions in DMEM supplemented with 10% fetal calf serum, 100 units/ml penicillin, 100 mg/ml streptomycin, and 2.5 mM glutamine. A875-NT (control) and A875 shp75 (knockdown) cell lines were described previously ^39^. Cell lysis and whole cell protein extraction were carried out in Lysis Buffer [50 mM Tris/HCl pH 7.5, 1 mM EDTA, 270 mM Sucrose, 1% (v/v) Triton X-100, 1 mM benzamidine, 1 mM PMSF, 0.1% (v/v) 2-mercaptoethanol, and in the presence of phosSTOP (Roche) phosphatase inhibitor cocktail mix as per manufacturer instructions]. Protein concentration was determined by Bradford Assay. Addition of Laemmli sample buffer and analysis by SDS-PAGE and Western Blot. Immunoblots were developed using the ECL Western Blotting Kit (Thermo Scientific) and exposed to Kodak X-Omat AR films.

For co-immunoprecipitation analysis of p75^NTR^/fascin interactions, A875 sh-p75^NTR^ and A875-WT cells were plated in a 10 cm tissue culture dish at a confluency of 3 × 10^6^ cells/dish in normal growth media. Twenty-four hours after plating, the cells were treated with 1μM 4A22 and NGF (100ng/ml) for 12 hr prior to harvest and lysis in lysis buffer (50 mM Tris/HCl pH 7.5, 1 mM EDTA, 270 mM Sucrose, 1% (v/v) Triton X-100, 0.1% (v/v) 2-mercaptoethanol and 60 mM n-Octyl-β-D glucopyranoside) containing protease inhibitor cocktail. The cellular extracts were then centrifuged at 4°C top speed on a benchtop centrifuge for 1 min and the supernatant was collected. For immunoprecipitation, A875 cell extracts were incubated for 16 hr at 4°C on a rotating wheel with 0.8μg of anti-p75^NTR^ antibody (ANT-007, Alomone) and then incubated with Sepharose protein-G beads (GE Healthcare). The beads were collected by brief centrifugation (2 min, 2500 rpm, 4°C), washed three times with 1ml of Wash Buffer (with composition same as the lysis buffer). After the last wash, pelleted beads were aspirated off the wash buffer followed by addition of Laemmli sample buffer and analysis by SDS-PAGE followed by Western Blot using anti-fascin antibody (MA5-11483, Invitrogen).

For detergent-insolubility assay of fascin, A875 NT cells were treated with or without 100 ng/ml NGF and 1µM of Np75-4A22 for 24h. A single cell suspension was obtained by trypsinization followed by centrifugation and detergent extraction with lysis buffer (with 0.5% Triton X-100). Then, immediately centrifuged at 8700 g for 3 min. The resulting pellets (containing the Triton X-100-resistant proteins) were suspended in lysis buffer without Triton X-100. In parallel, cells were centrifuged and then directly suspended in lysis buffer. The Triton X-100-resistant proteins and total cell lysates were analyzed by SDS–PAGE and Western blotting with anti-fascin antibody.

### Cell viability, motility and chemotaxis analyses

To measure the cytotoxic effect of Div17E5, A875 NT or shp75 cells were seeded at 2,000 cells per well (in 100 μL DMEM) in a 96-well plate and incubated for 12h. Next, Div17E5 at various concentrations was added and incubated at 37°C in 5% CO2 for 2 days. Following treatment, 10μL Presto Blue® (resazurin, ThermoFIsher) was added to each well of the 96-well plate and incubated for 2 hours at 37°C in 5% CO2. Cellular fluorescence in each well was measured at 535 nm excitation and 615 nm emission using a microtiter plate reader. Data was converted into the relative cell viability (%) from the fluorescence of cells in each treatment relative to that of the DMSO control group (set as 100%). The student’s t-test was used to compare data between cell lines. For wound healing assay, cells were grown until confluency and subjected to a wound across the cell mono-layer. Cells were then incubated with serum-free media either with DMSO or a small molecule (with or without 5µM Q_VD_poH) and imaged at specific time-points. Cell migration is reflected as fold change in wound area over time.

For Boyden chamber assay, A875 NT cells were serum-starved for 16Hrs and then introduced in serum-free media in 4 different conditions, DMSO, 5µM Q_VD_poH, a small molecule and 5µM Q_VD_poH + small molecule to the top chamber of the Matrigel coated trans-well chamber. Two different chemoattractant, 50ng/ml NGF or 10% FBS were added to the bottom chamber. Assays were carried out for 8Hrs followed by fixation, staining and imaging. Number of cells that crossed the chamber were counted as invading cells.

## Synthetic methods

### Procedure A (Amination 1)

2,6-Dichloropyrazine (1-2 eq) and the substituted aniline (1-2 eq) were dissolved in anhydrous N,N-dimethylformamide (DMF) and cooled to -78 °C. Sodium tert-butoxide (NaO-tBu) 2.5 M in tetrahydrofuran (THF) or sodium tert-pentoxide (NaO-t-pent) 40 % in THF (3.5 eq) was added dropwise and the reaction mixture stirred and allowed to warm to room temperature. Stirring was continued for between 3 to 24 hours, by monitoring the reaction via TLC and/or LCMS. On completion, the reaction mixture was dissolved in dichloromethane (DCM), washed twice with water and once with saturated brine solution. The organic portion was dried with magnesium sulfate (MgSO4), filtered and concentrated, before being purified by column chromatography on a SNAP (Biotage) automated purification system using isohexane: ethyl acetate (0-100 %).

### Procedure B (Amination 2)

This method was used for some compounds for which Procedure A failed to yield the product of interest. 2,6-Dichloropyrazine (1-2 eq) and the substituted secondary amine (1-2 eq) were dissolved in anhydrous acetonitrile at room temperature. N, N-Diisopropylethylamine (DIPEA, 2.5 eq) was added to the reaction mixture and stirred at 80 °C for 3-4 hours. The reaction mixture was cooled to room temperature, dissolved in DCM, washed twice with water and once with saturated brine solution. The organic portion was dried with MgSO4, filtered and concentrated, before being purified by column chromatography on SNAP (Biotage) automated purification system using DCM (100 %).

### Procedure C (Suzuki coupling 1)

Substituted 6-chloro-pyrazine-2-amine (1-2 eq) (obtained for Procedure A), substituted phenylboronic acid (1-2 eq), K2CO3 (3 eq), and [1,1′-bis(diphenylphosphino)ferrocene]dichloropalladium(II), complex with dichloromethane (Pd(dppf)Cl2:DCM, 0.1 eq) were dissolved in 1,4-dioxane: H2O (2:1 mixture) in a microwave tube. Microwave irradiation was carried out at 120 °C for 12 min. After LCMS showed complete conversion to product of interest, the reaction mixture was dissolved in DCM, washed twice with water and once with saturated brine solution. The organic portion was dried with MgSO4, filtered and concentrated, before being purified by column chromatography on SNAP (Biotage) automated purification system using isohexane: ethyl acetate (0-100 %).

### Procedure D (Suzuki coupling 2)

This method was used for some compounds for which Procedure C failed to yield the product of interest. 2,6-Dichloropyrazine (1-2 eq), substituted phenylboronic acid (1-2 eq), K2CO3(aq) or Na2CO3(aq) (2.0 M, 2-3 eq), and tetrakis(triphenylphosphine)-palladium(0) (Pd(PPh3)4, 0.05 eq) were dissolved in toluene: ethanol (2:1 mixture). The reaction was carried out using either one of the two following procedures:

(i) In a microwave tube, microwave irradiation was carried out at 70 °C for 30 min. After LCMS showed complete conversion to product of interest, the reaction mixture was dissolved in DCM, washed twice with water and once with saturated brine solution. The organic portion was dried with MgSO4, filtered and concentrated, before being purified by column chromatography on SNAP (Biotage) automated purification system using isohexane: ethyl acetate (0-100 %).
(ii) In a round-bottom flask, the reaction mixture was refluxed at 120 °C for 7 h. After LCMS showed complete conversion to product of interest, the solvent was removed under vacuum. The crude product was dissolved in DCM, washed twice with water and once with saturated brine solution. The organic portion was dried with MgSO4, filtered and concentrated, before being purified by column chromatography on SNAP (Biotage) automated purification system using isohexane: ethyl acetate (0-100 %).

### Pharmacokinetics and toxicology

Plasma pharmacokinetics of Np75-4A22 was investigated following single intravenous (3 mg/kg) or oral (10 mg/kg) dose administrations in C57BL/6 mice of both sexes. Blood samples were collected from the retro orbital plexus at pre-dose, 0.08, 0.25, 0.5, 1, 2, 5, 10 and 24 hs (for intravenous) and pre-dose, 0.25, 0.5, 1, 2, 4, 5, 10 and 24 hs (for oral). Toxicity of Np75-4A22 was tested in a 3-week repeated dose study in C57BL/6 mice following oral gavage administration. Pharmacokinetics and toxicology studies were carried out by Sai Life Sciences Limited, Hyderabad, India.

### Melanoma lung metastasis assay, immunohistochemistry and immunofluorescence

Eight-week-old female SCID mice (Charles River Laboratories, Shrewsbury, MA, USA) were housed in groups of three to five and maintained on a 12-h light/dark schedule with a temperature of 22 °C±1 °C and a relative humidity of 50%±5%. Food and water were available ad libitum. All procedures were reviewed and approved by the University of Calgary Animal Care Committee. A875-GFPLuc (A875-NT) or A875-GFPLuc shp75 (A875-shp75) cells harvested using Puck’s EDTA were resuspended in PBS and injected intravenously into SCID mice (5 x10^5^ cells/200 μl per mouse). Np75-4A22 was dissolved in 0.5% sodium carboxymethyl cellulose (NaCMC) in reverse osmosis H_2_0 and administered by oral gavage at 200 mg/kg/day starting one day prior to intravenous injection of tumor cells and continued for 3 one-week cycles (cycle: 5 days on drug, 2 days off drug). Metastatic spread to the lungs was monitored weekly by luciferase activity using the Xenogen IVIS-200 Optical *in vivo* luciferase imaging system as described previously ^56^ and by immunohistochemistry (IHC) at designated time points. IHC was performed after formalin fixation, paraffin embedding and sectioning of the lungs, using a human-specific mouse polyclonal nucleolin antibody (1:500 dilution) and detected using Dako liquid DAB+ Substrate Chromogen System (Dako: k3468, California, US) followed by a hematoxylin counterstain (Sigma, Oakville, ON, Canada). Sections were mounted and imaged on an Olympus phase contrast microscope (Olympus Life Science, Tokyo, Japan). For immunofluorescence, sections were stained with anti-human nucleolin and anti-human p75^NTR^ antibodies (1:1000), subsequently detected with 1:200 dilutions of Alexa Fluor™ 488 Goat anti-Mouse IgG (H+L) Highly Cross-Adsorbed Secondary Antibody or Alexa Fluor™ 568 Goat anti-Rabbit IgG (H+L) Cross-Adsorbed Secondary Antibody. Sections were further stained with DAPI to detect all nuclei. Between each treatment, TBS Automation Wash Buffer (Biocare Medical TWB945M) was employed. Images were taken with LSM800 confocal microscope and ZEN system.

### Statistical analysis

Statistical Analysis Software (SAS Institute, Inc.) and GraphPad Prism (versions 4 or 8; GraphPad Software Inc, San Diego, CA, USA) were used for statistical analyses. Experimental data was collected from multiple experiments and reported as the treatment mean ± SEM. Statistical significance was calculated using the Student *t*-test, Mann Whitney or one-way ANOVA. *p* value of less than 0.05 was considered statistically significant.

## Acknowledgements

We thank Ket Yin Goh for technical assistance. This research was funded by grants to C. F. I. from the Singapore National Medical Research Council (NMRC/OFIRG/MOH-000224), Swedish Research Council (2020-01923) and Swedish Cancer Society (18-0670 and 18-0829) and to D.L.S. from the Canadian Cancer Society and Robert C. Westbury Endowment.

## Author contributions

V.L.-R. performed the majority of cell culture experiments, screening and validation of all Div17E5 derivatives and compiled a preliminary draft of the manuscript and figures;

S.A.N. performed all medicinal chemistry and SAR studies resulting in compounds of Series 3 and 4 including the lead compound Np75-4A22; X.L. and J.Z. performed the metastasis studies in SCID mice; A.R. performed co-immunoprecipitation studies; S.S. performed the initial screening that identified Div17E5; M.S. performed preliminary medicinal chemistry studies resulting in compounds of series 1 and 2; D.L.S. designed and supervised all cancer studies; C.F.I. C.F.I. conceived the original project, contributed to project coordination and data analysis, and wrote the manuscript.

## Supplementary text

### Analysis of structure-activity relationships in Series 3 and 4

MS_18 and MS_63 were modified to probe for the effects of i) mono vs. multiple substitutions; (ii) positional isomers; (iii) bio-isosteric replacement, e.g., O to S; (iv) electron-withdrawing vs. -donating groups; (v) bulkier vs. smaller groups; and (vi) amino vs. amide linker, to yield compounds in Series3A, 3B, 4A and 4B.

For 3A analogs, we found that: i) 4-methoxy and 4-methyl were preferred over the 2- or 3-regio isomeric substituents (MS_18 more potent than Np75-3A01, -3A02, and Np75-3A21 better than -3A19 and -3A20); ii) when 4-methoxy (4-OCH3) was replaced with the isosteric 4-methylthio (4-SCH3), or 4-methylsulfonyl (4-SO2CH3) groups, activity was lost; iii) multiple electron-donating methoxy substitutions, as well as di- and trimethyl groups were not favored; iv) electron-withdrawing groups such as nitro (NO2), cyano (CN), trifluoromethyl (CF3), carboxylic acid (COOH), and halogen motifs were also unfavorable; v) bulkier pentyloxy, phenoxy and benzyloxy groups did not also improve activity; vi) when the amine linker was converted to an amide in Np75-3A49 and Np75-3A51, activity was lost. Np75-3A21 was identified as the best hit from this series.

For 3B analogs: i) none of the modifications employed improved the activity of MS_18; ii) the best compound in this series was Np75-3B03, which possessed the same 4-bromo-3-chloro substitution found in Div17E5 but still less potent than MS_18.

For 4A analogs: i) unlike Series 3A, multiple electron-donating methoxy substitutions (specifically 3,4-dimethoxy), as well as dimethyl groups (i.e., 2,3-dimethyl) were favored in this series; ii) electron-withdrawing and bulkier phenoxy and benzyloxy motifs were also not preferred in this series. The best compound in the 4A series was Np75-4A22.

For 4B analogs, none of the modifications employed improved the activity of MS_63.

**Figure S1.**
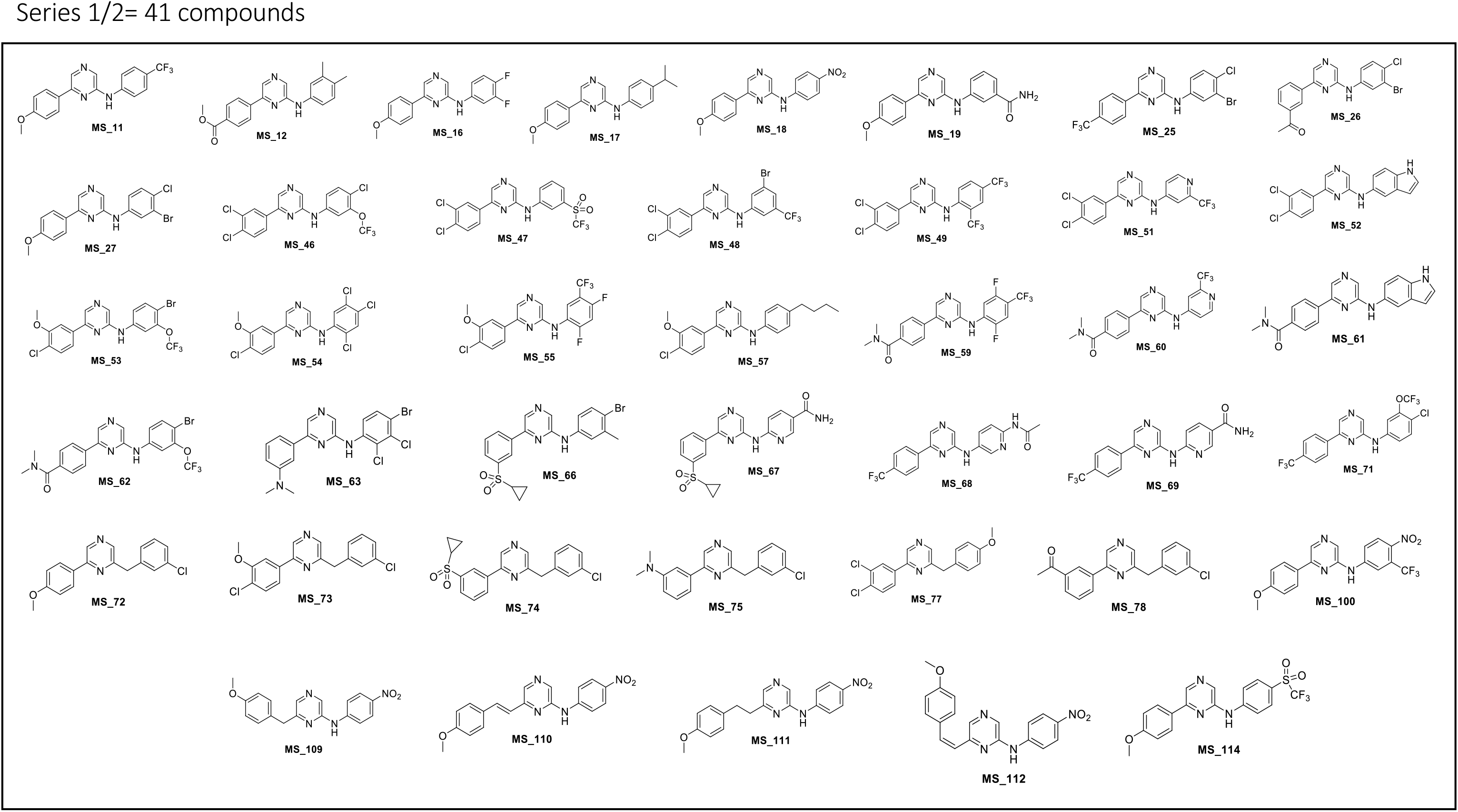
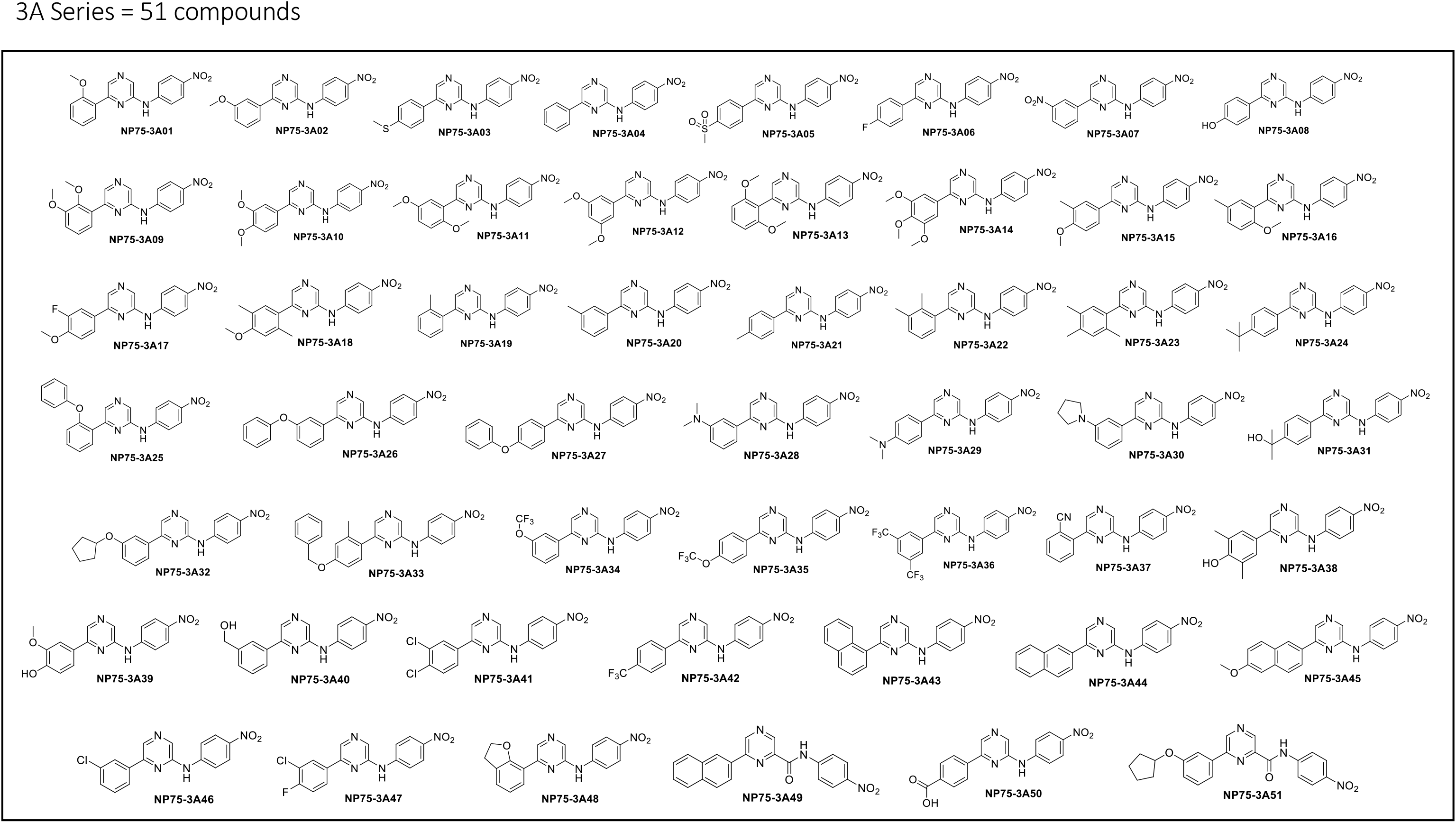
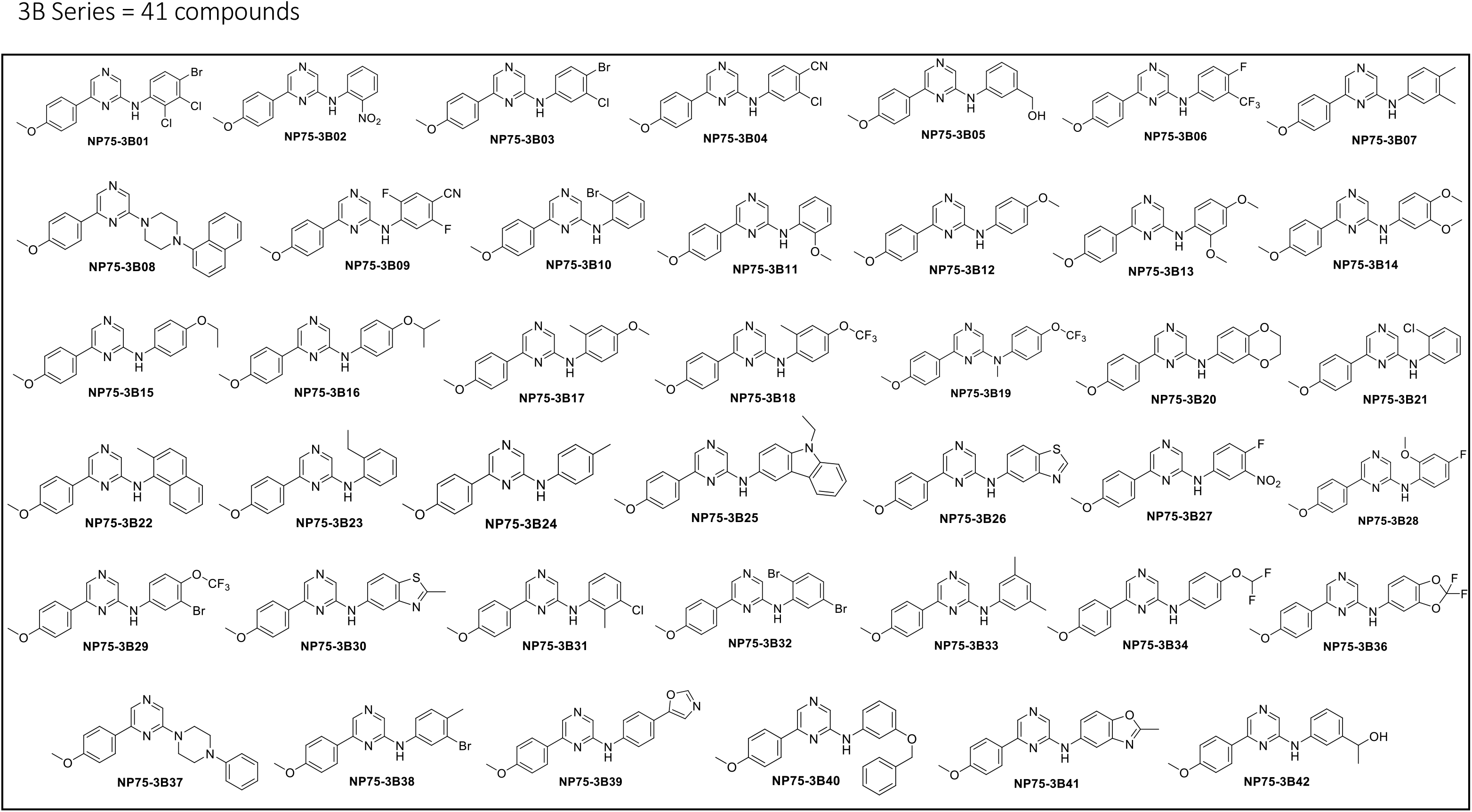
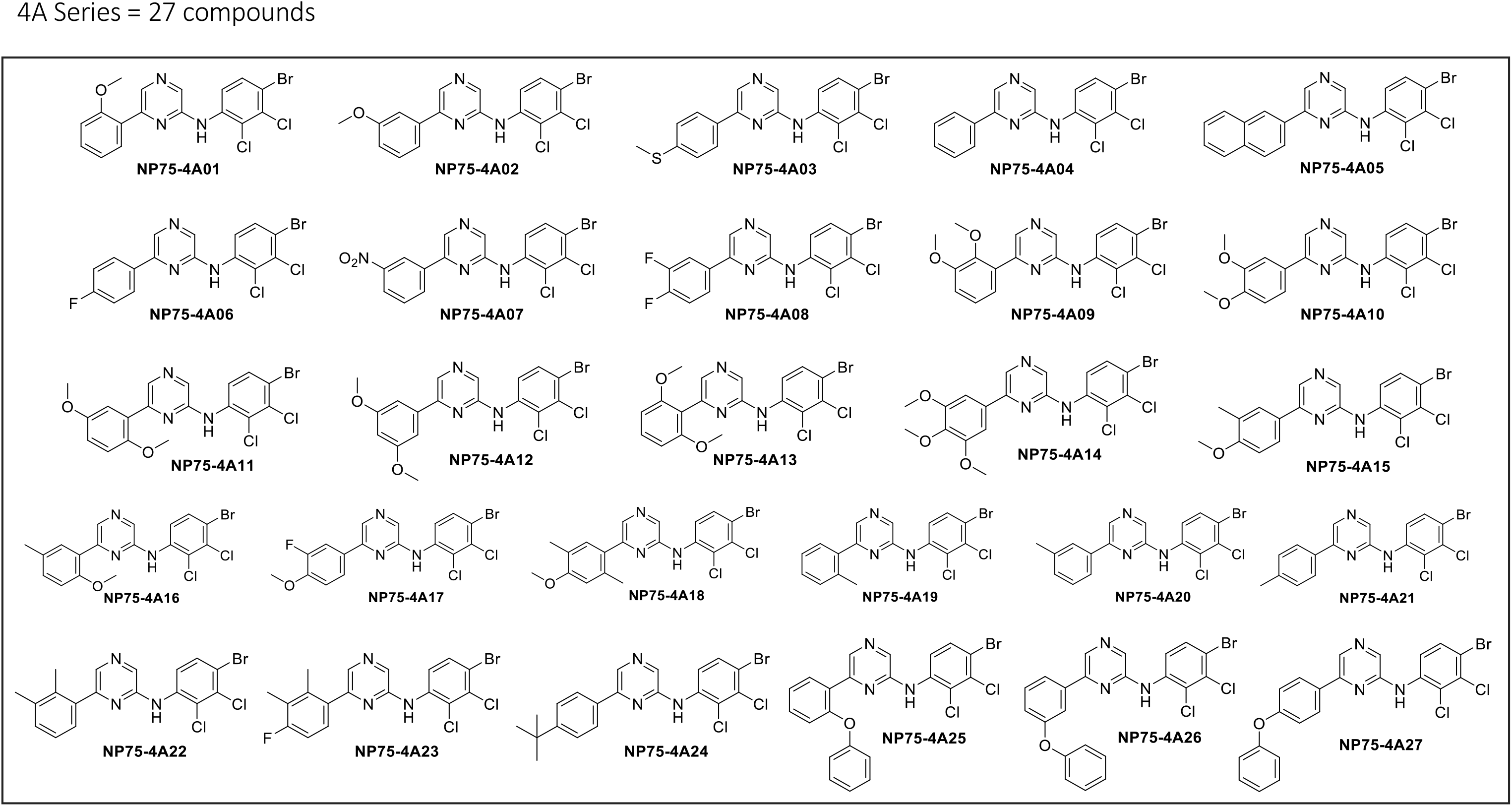
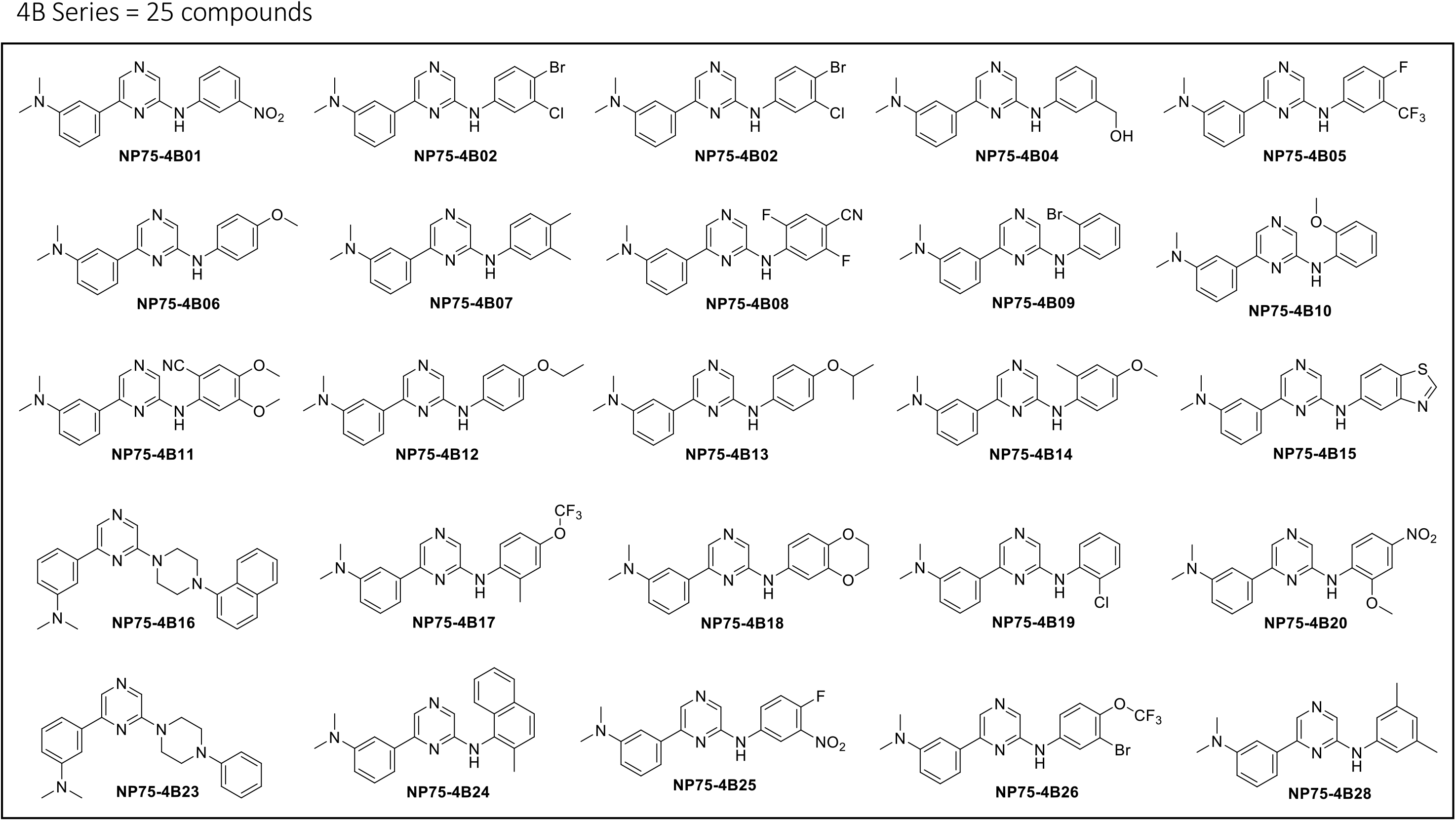
Chemical structures of Div17E5 analogs. (A) Structures compounds in Series 1 and 2 (A), Series 3A (B), Series 3B (C), Series 4A (D) and Series 4B (E).

**Figure S2.**
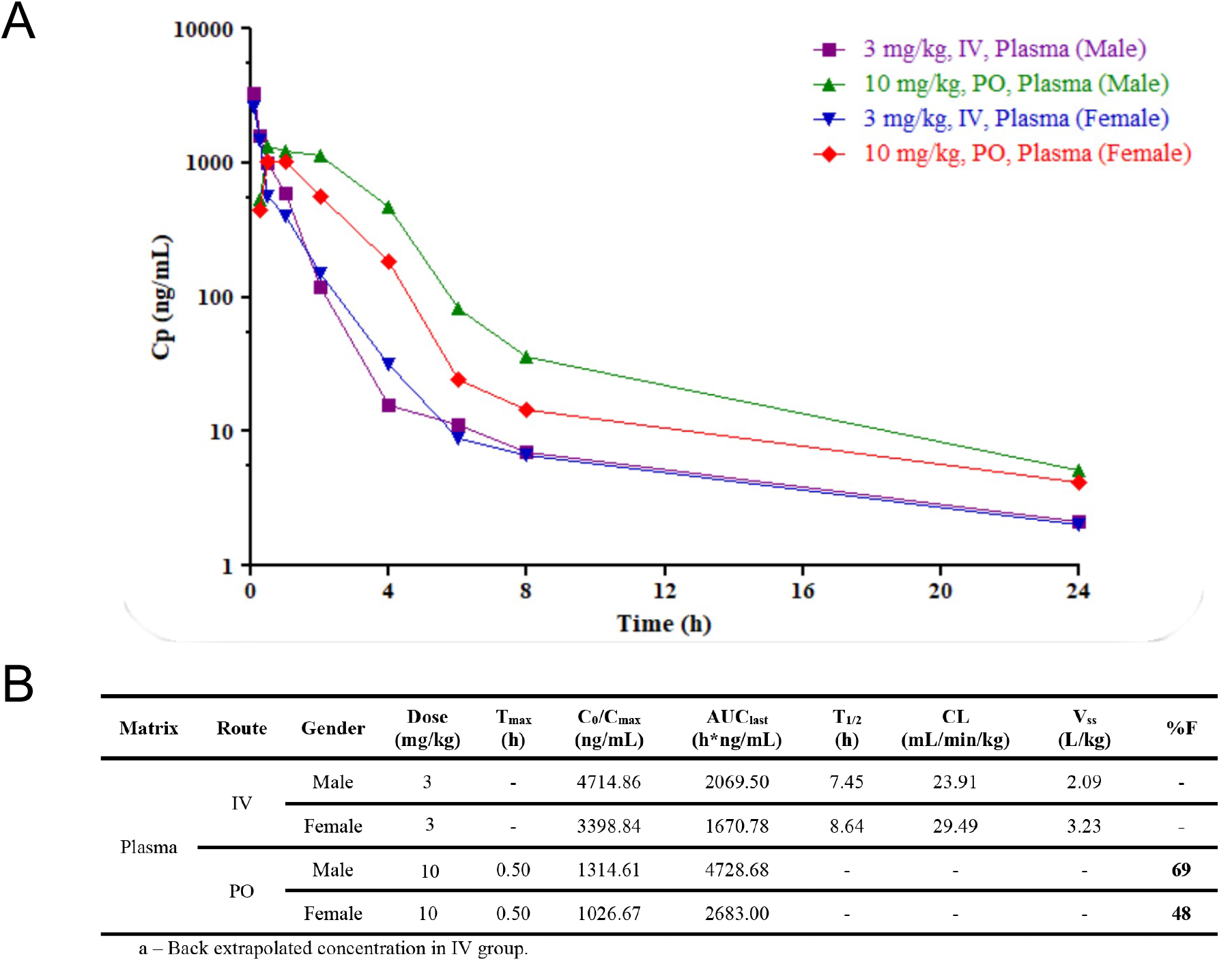
Pharmacokinetic study of Np75-4A22 in mice. (A) Graph shows plasma concentrations of Np75-4A22 in male and female C57BL/6 mice (N=9) after a single intravenous (3mg/kg) or oral (10mg/kg) administration of 10mg/kg. Plasma was collected at pre-dose, 0.25, 0.5, 1, 2, 4, 5, 10 and 24 hs and assessed for Np75-4A22 using LC/MS/MS. (B) Pharmacokinetic parameters of Np75-4A22 in plasma following a single intravenous (3mg/kg) or oral (10mg/kg) in male and female C57BL/6 mice. Cmax: maximum (peak) plasma drug concentration (amount/volume). Tmax: time to reach maximum (peak) plasma concentration following drug administration. AUClast: area under the plasma concentration-time curve from time zero to time of last measurable concentration (time/volume). Bioavailability (%F): extent and rate at which the active moiety (drug or metabolite) enters systemic circulation.

**Figure S3.**
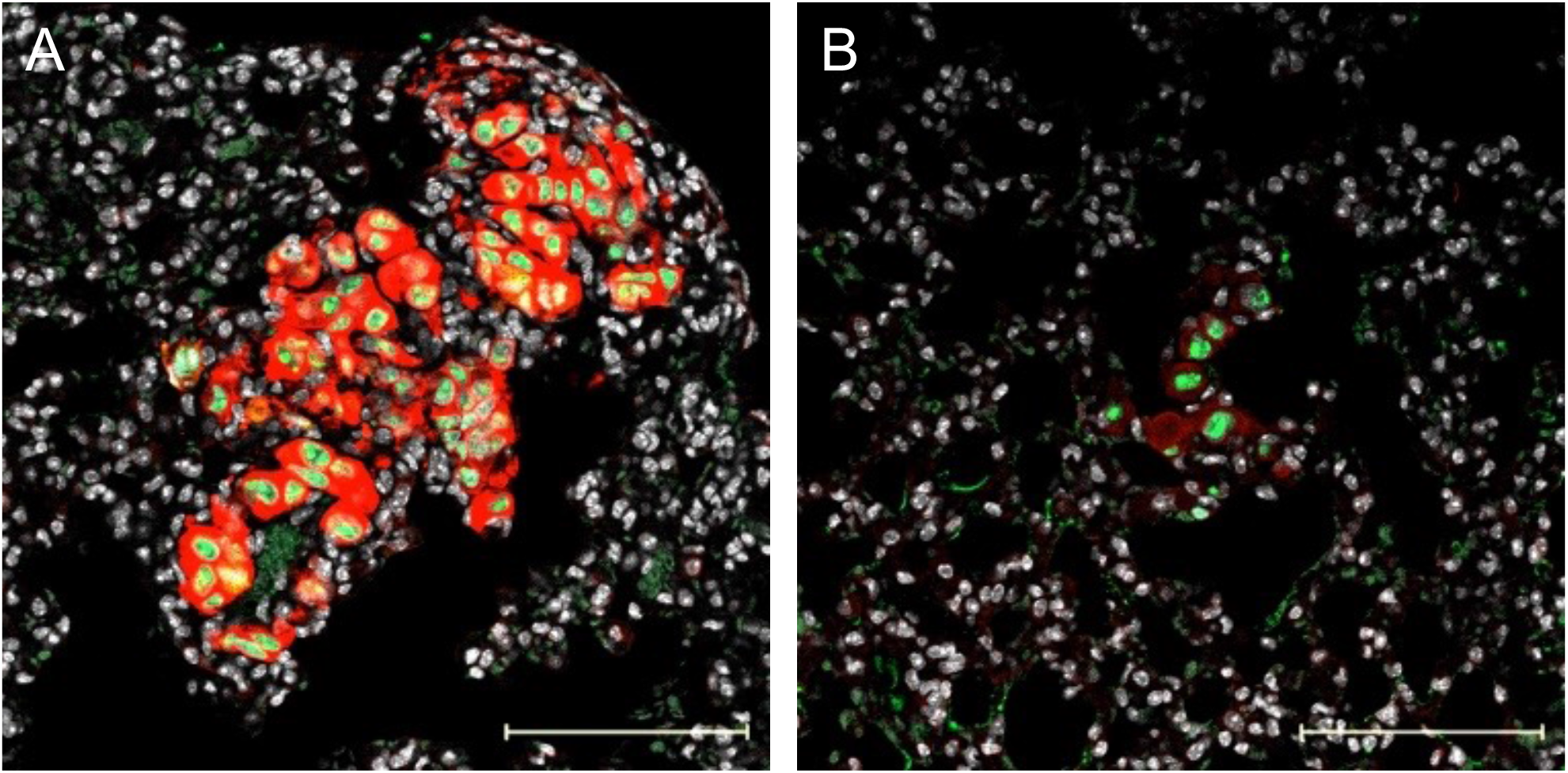
Histological analysis of p75NTR expression in lung metastasis produced by A875 cells. p75^NTR^ immunostaining (red) in lung metastasis induced by A875-NT (A) or shp75-A875 (B) cells counter-stained for human nucleolin (green) and DAPI (white). Scale bar, 100μM.

